# Differential roles for the spindle assembly checkpoint in error surveillance and mitotic timing

**DOI:** 10.1101/2025.06.14.659703

**Authors:** Ryan K. Kim, Kuan-Chung Su, Mary-Jane Tsang, Nolan Maier, Brittania Moodie, Iain M. Cheeseman

**Affiliations:** Whitehead Institute for Biomedical Research, Cambridge, United States; Department of Biology, Massachusetts Institute of Technology, Cambridge, United States; Peter MacCallum Cancer Centre, Melbourne, Victoria 3000, Australia; Department of Microbiology, Harvard Medical School, Boston, MA, United States

## Abstract

The Spindle Assembly Checkpoint (SAC) plays critical roles in regulating mitotic fidelity and progression. Here, we utilized a SAC-deficient cell line lacking the full-length Cdc20 translational protein isoform (Cdc20 ΔFL) to define its differential genetic interactions using CRISPR/Cas9-based gene targeting. Cdc20 ΔFL cells display synthetic lethal relationships with gene targets required for proper chromosome segregation, highlighting the critical role of the SAC in error surveillance. Surprisingly, we found that the checkpoint component Mad2 becomes dispensable for viability in Cdc20 ΔFL cells. Prior work suggested that Mad2 acts as an essential mitotic “timer” to control mitotic duration in unperturbed cells. Instead, our functional analysis indicates that the mitotic timer depends on the interdependent and overlapping actions of: (1) Mad2 inhibition of APC/C-Cdc20, (2) Cdk1-mediated phosphorylation of Cdc20, and (3) total Cdc20 protein levels. Simultaneously perturbing these pathways results in near immediate mitotic exit and catastrophic chromosome mis-segregation.

## Introduction

Cell cycle checkpoints act to delay or halt cell cycle progression in the presence of errors. Cell cycle checkpoints were first defined by the identification of the Rad9-dependent DNA Damage Checkpoint in budding yeast ^1^ and subsequently by the identification of the Spindle Assembly Checkpoint ^2, 3^. In yeast, these checkpoint factors are dispensable for viability under normal growth conditions but become critically needed when the cells experience DNA damage or mitotic errors. The checkpoint was originally defined as a surveillance mechanism that is only “activated” in the presence of errors, allowing cells to correct damage before the cell cycle proceeds ^1^. However, subsequent studies in vertebrate cells instead proposed that the Spindle Assembly Checkpoint acts in all cells to prevent premature mitotic progression and is only “satisfied” (inactivated) once correct chromosome alignment is achieved ^4^. This model is based in part on studies in mammalian cell and mouse models that identified essential requirements for checkpoint proteins (MAD2L1, MAD1L1, BUB1, BUB1B, TTK). For example, even in the absence of mitotic errors, loss of the checkpoint protein Mad2 (MAD2L1) leads to a dramatically accelerated mitosis and severe chromosome mis-segregation ^5, 6^. Due to these phenotypes, Spindle Assembly Checkpoint components have also been proposed to act as the timer for mitotic duration ^5^.

Our recent work generated a human cell line (referred to here as Cdc20 ΔFL) that lacks the longer translational protein isoform of Cdc20, a key downstream target of the Spindle Assembly Checkpoint ^7^. Cdc20 ΔFL cells are viable and progress through mitosis normally under unperturbed conditions. However, due to the absence of the Cdc20 N-terminal checkpoint interaction motifs ^8–10^ in the alternative truncated Cdc20 isoforms, Cdc20 ΔFL cells are unable to arrest in mitosis in the presence of anti-mitotic drugs ^7^. This ability of the Cdc20 ΔFL mutants to remain viable despite a defective checkpoint suggests that the mammalian Spindle Assembly Checkpoint acts similarly to the original observations in budding yeast. In this case, the essential roles and strong phenotypes associated with the loss of other checkpoint proteins could suggest that these proteins play dual roles in both the surveillance mechanism of the checkpoint and the regulation of mitotic timing or chromosome segregation. For example, Mps1 (TTK) phosphorylates multiple kinetochore substrates to regulate kinetochore-microtubule attachments and kinetochore function ^11^, but also plays a role in preventing premature mitotic progression ^12^. Similarly, the dramatic acceleration of mitotic duration in cells lacking Mad2 suggests that Mad2 acts as a timer to control mitotic duration to ensure that chromosomes have sufficient time to align at the metaphase plate. This is in addition to its surveillance role of monitoring mitotic fidelity ^3^. However, based on these differing functional requirements, the relative contributions of the checkpoint as an error surveillance pathway vs. a signaling pathway that must be “satisfied” to control mitotic timing remain elusive.

Here, we leverage the behavior of Cdc20 ΔFL cells to provide a unique genetic background to explore the functional contributions of the Spindle Assembly Checkpoint in human cells. Using CRISPR/Cas9-based functional genetics, we demonstrate that Cdc20 ΔFL cells display a synthetic lethal relationship with gene targets whose loss results in increased chromosome segregation defects, consistent with the classically defined role for the Spindle Assembly Checkpoint. Surprisingly, despite the essential roles for checkpoint components in control cells, we demonstrate that multiple checkpoint factors, such as Mad2, become dispensable for viability in the Cdc20 ΔFL background. By analyzing these behaviors, we reveal that the basis for the “timer” for mitotic progression extends beyond the action of Mad2 in restraining premature APC/C activation. Our work demonstrates that this timer is regulated by the interdependent and overlapping actions of: (1) Mad2 inhibition of Cdc20-bound APC/C, (2) CDK-dependent phosphorylation of Cdc20 to block its activity, and (3) the total levels of Cdc20 available for APC/C activation. By perturbing these pathways individually and in combination, we were able to eliminate the mitotic timer, resulting in near immediate mitotic exit and catastrophic chromosome mis-segregation.

## Results

### Cdc20 **Δ**FL cells are sensitized to errors in chromosome alignment and spindle assembly

Cervical cancer-derived HeLa cells ^7^, leukemia-derived K562 cells (this study), and lung adenocarcinoma-derived A549 cells (this study) lacking the full length Cdc20 translational isoform are viable (Fig. 1A-B, S1A-F; referred to here as Cdc20 ΔFL). However, Cdc20 ΔFL cells are unable to undergo a mitotic arrest when treated with drugs that disrupt chromosome segregation ^7^. The absence of a functional Spindle Assembly Checkpoint provides an ideal genetic background to identify additional gene targets that modulate mitotic fidelity and progression ^13–15^. We therefore used CRISPR/Cas9-based functional genetics to compare the growth requirements for control and Cdc20 ΔFL cells (Fig. 1C,D). To conduct these large-scale analyses, we introduced a lentiviral library targeting 1411 genes enriched for factors related to microtubule function/spindle assembly, cell cycle control, checkpoint regulation, kinetochore function, and chromosome organization/maintenance ^16^ (Fig. 1C). We conducted these screens in Cdc20 ΔFL mutants for both HeLa cells (Fig. 1D, Supplementary Table 1) and K562 cells (Fig. S1G, Supplementary Table 1). Following lentiviral transduction and selection, we assessed the change in sgRNA abundance over 14 population doublings. We then calculated a CRISPR score that corresponds to the average log_2_-fold change in the single-guide abundance, with a score of -1 or lower corresponding to genes that are essential for cellular viability and a score of 0 corresponding to genes whose knockouts have minimal effects on cellular fitness. In these screens, most genes displayed similar growth effects in both control cells and Cdc20 ΔFL cells, with an R^2^ value of 0.8255 (Fig. 1D) in HeLa cells and 0.8160 in K562 cells (Fig. S1G). However, we observed cases in which Cas9-mediated gene targeting resulted in differential effects between control and Cdc20 ΔFL cells. For example, for some genes with roles in chromosome alignment/spindle assembly (CENPE, MKI67, KIF2C) or cell cycle control (FZR1, CCNB2), the negative growth defects caused by their knockout were further exacerbated in the HeLa Cdc20 ΔFL background.

**Fig. 1:**
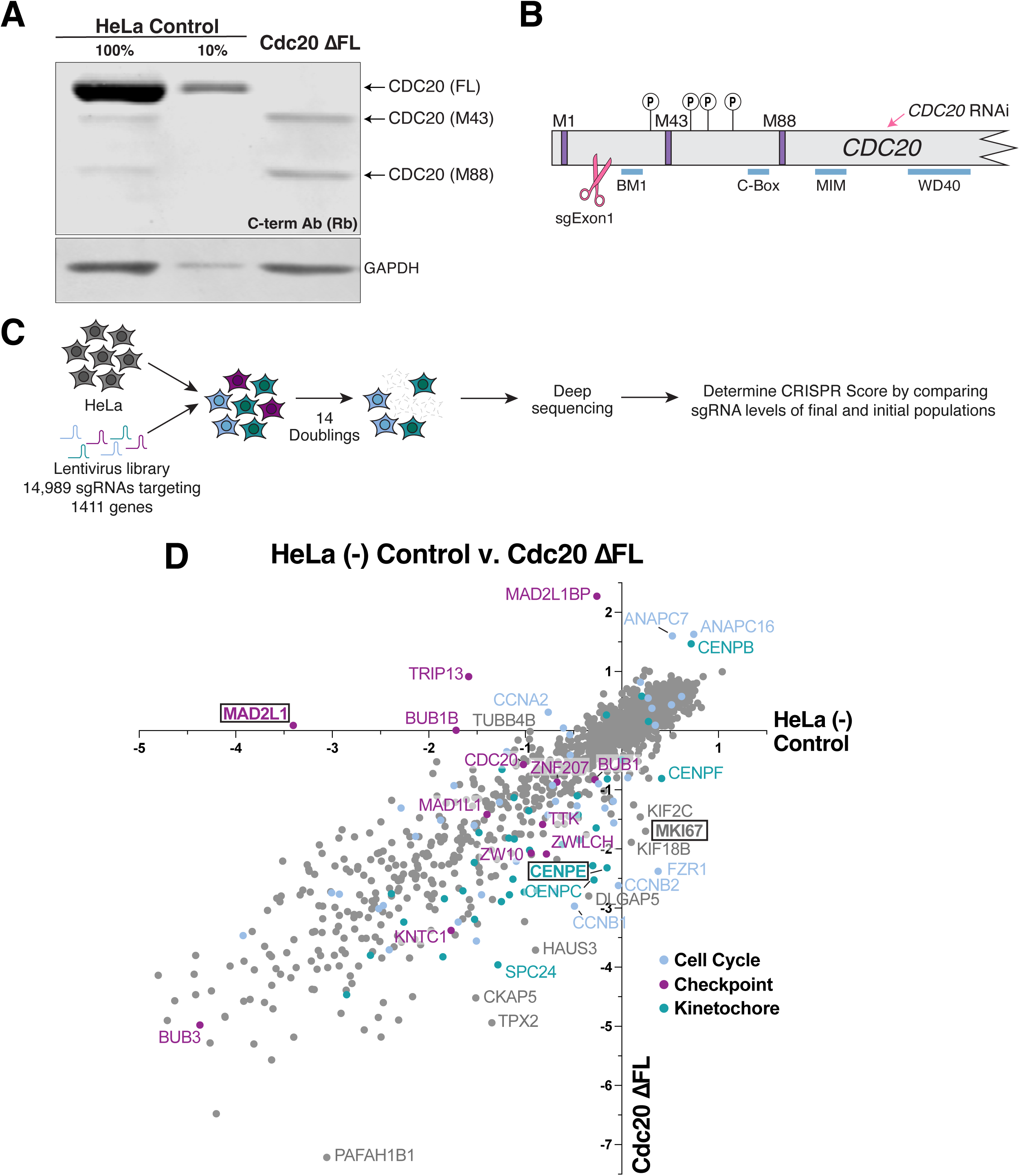
Functional genetics screen reveals differential modulators of mitotic fidelity and progression in Cdc20 ΔFL cells compared to control cells. a, Western blot analysis of endogenous Cdc20 in HeLa and Cdc20 ΔFL (Clone #3) cells using antibodies recognizing the C-terminus of human Cdc20 (aa 450-499). GAPDH was used as a loading control. b, The *CDC20* open reading frame, indicating the targeting of *CDC20* RNAi and sgExon1. The *CDC20* Met1, Met43, and Met88 start sites, and relevant N-terminal CDK phosphorylation sites are labeled. c, Schematic showing the workflow for the functional genetics screen. d, Scatter plot showing the CRISPR scores in HeLa (-) control cells vs. Cdc20 ΔFL cells (Clone #1).

To evaluate these findings further, we focused on gene knockouts that had minimal effects on cellular viability on their own but were lethal in the Cdc20 ΔFL background. We first used a fluorescence-based co-culture competition assay (Fig. 2A) to evaluate the kinetochore-localized kinesin CENPE. This assay monitors the relative growth behavior of two cell lines, allowing us to make sensitized and direct comparisons of their proliferation. In pairwise assays, we found that ΔCENPE + Cdc20 ΔFL cells displayed a substantial growth deficit compared to ΔCENPE single knockouts alone (Fig. 2B). In addition, based on immunofluorescence analysis, we found that ΔCENPE + Cdc20 ΔFL double mutant cells displayed a modest but reproducible increase in anaphase errors, as well as increased micronuclei formation compared to ΔCENPE cells (Fig. 2C, D, S2A, B). Similarly, a gene knockout of MKI67, a chromosomal periphery protein and surfactant ^17, 18^, had more severe fitness defects in Cdc20 ΔFL cells than in control HeLa cells (Fig. 2E). ΔMKI67 + Cdc20 ΔFL cells reproducibly displayed a slight increase in anaphase errors (Fig. 2D) and micronuclei formation (Fig. S2B). The anaphase chromosome segregation errors observed in CENPE and MKI67 knockout cells are predicted to be prevented by the SAC, which delays the onset of anaphase in the presence of defects to allow cells to resolve these errors prior to chromosome segregation. However, in the SAC-defective Cdc20 ΔFL cells, these errors would go unchecked and remain unresolved, eventually resulting in an accumulation of genomic instability and cell death. Together, this functional genetic approach reveals important contributions of the SAC in surveilling mitotic errors and ensuring proper chromosome segregation.

**Fig. 2:**
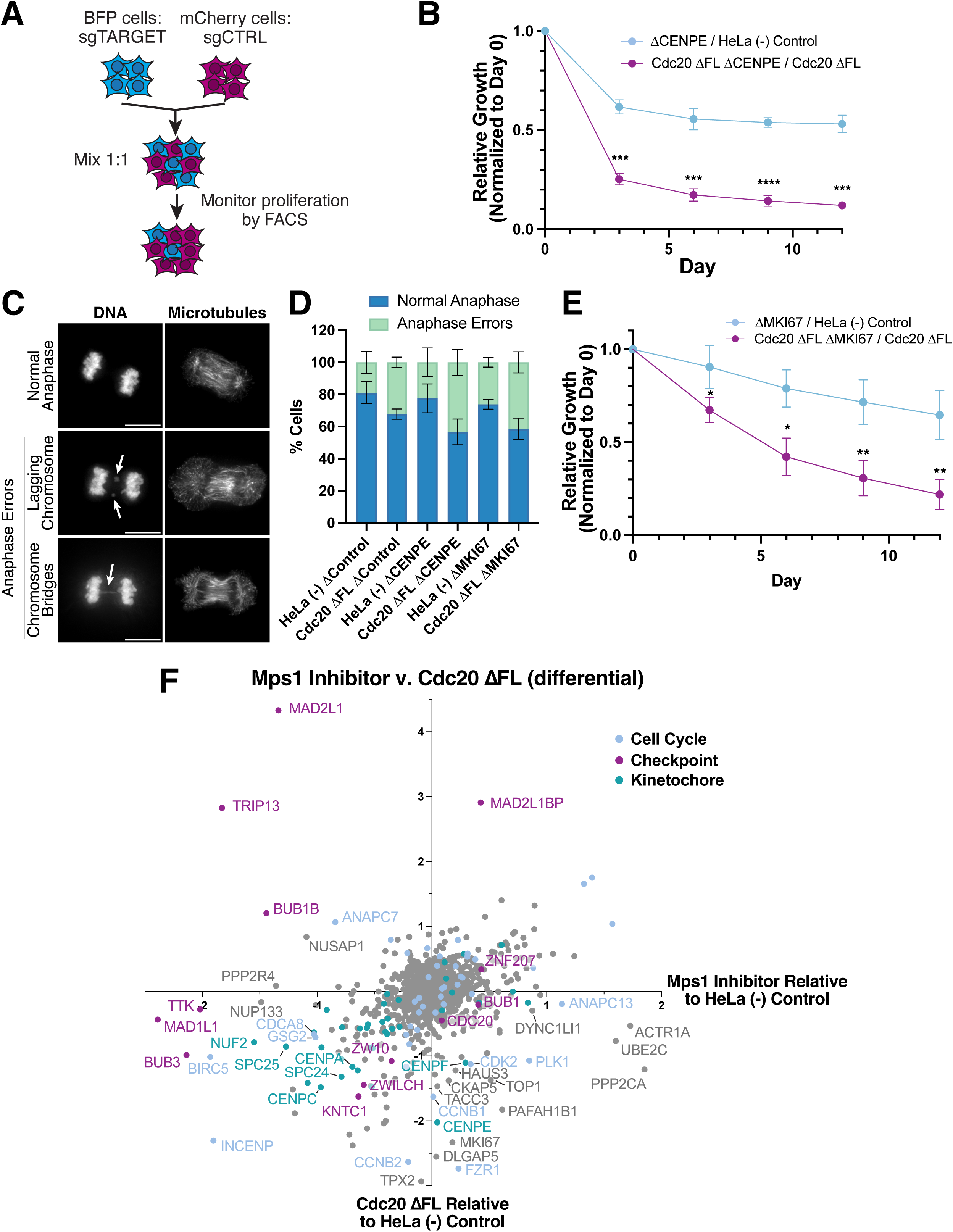
Cdc20 ΔFL cells display a synthetic lethal relationship with a CENPE knockout or a MKI67 knockout. **a,** Schematic showing the workflow for the fluorescence-based competition assay. **b,** Competitive growth assay to test the relative fitness of inducibly eliminating CENPE in HeLa control or Cdc20 ΔFL cells (Clone #1). Data points reflect mean ± s.e.m.; *n* = 3 experimental replicates. **c**, Representative deconvolved Z-projected immunofluorescence images of anaphase cells from HeLa and Cdc20 ΔFL cells in which CENPE or MKI67 is inducibly eliminated. Images show microtubules (DM1α) and DNA (Hoeschst). Arrowheads indicate the specified anaphase error. Scale bars, 10 µM. **d,** Percentage of cells with anaphase defects in HeLa and Cdc20 ΔFL (Clone #1) cells in which a control locus (ΔHS1), CENPE, or MKI67 is inducibly eliminated. *n* > 400 cells per cell line, across four experimental replicates. Mean percentage is shown, error bars indicate SD. **e,** Competitive growth assay to test the relative fitness of inducibly eliminating MKI67 in HeLa control or Cdc20 ΔFL cells (Clone #1). Data points reflect mean ± s.e.m.; *n* = 3 experimental replicates. **f,** Scatter plot showing the differential CRISPR scores in Mps1 inhibitor treated vs. Cdc20 ΔFL cells (Clone #1). The differential was calculated relative to HeLa (-) control cells for both Mps1 inhibitor treated cells and Cdc20 ΔFL cells. For **b,e**, Student’s two-sample-*t-*tests with two-tailed distribution comparing relative ratios shown for Days 3-12. \**P*<0.05, \*\**P*<0.01, \*\*\**P*<0.001, \*\*\*\**P*<0.0001.

### Cdc20 **Δ**FL cells and Mps1 inhibitor-treated cells display distinct genetic dependencies

As an alternative strategy to test synthetic effects in cells with a compromised Spindle Assembly Checkpoint (SAC), we also conducted CRISPR/Cas9-based screening in the presence of AZ-3146, a small molecule inhibitor targeting the checkpoint Mps1 kinase (Fig. S2D, Supplementary Table 1). We then compared the relative CRISPR scores of gene targets in Cdc20 ΔFL and Mps1 inhibitor-treated backgrounds to assess their differential effects relative to control cells (Fig. 2F, Supplementary Table 1). Both Cdc20 ΔFL and Mps1 inhibitor-treated cells displayed increased sensitivity for knockouts of kinetochore genes (CENPA, CENPC, SPC24, SPC25, NUF2, etc.) and Chromosomal Passenger Complex components and related factors (INCENP, BIRC5, CDCA8, GSG2/Haspin), consistent with a role for the SAC surveillance pathway in protecting cells from chromosome mis-segregation. However, we also identified gene targets that displayed differential behaviors between these conditions. Knockouts of subunits of the Anaphase Promoting Complex and ubiquitin-related machinery (ANAPC13, UBE2C), subunits of the dynein-dynactin complex (DYNC1LI1, ACTR1A), and of protein phosphatase 2 (PPP2CA) displayed appreciably improved growth in Mps1 inhibitor-treated cells, but not in Cdc20 ΔFL cells. Reciprocally, knockouts of the cyclins CCNB1 or CCNB2, or of APC/C adapter FZR1 displayed reduced growth in Cdc20 ΔFL cells, but not in Mps1 inhibitor-treated cells. Finally, as described below, the Cdc20 ΔFL mutant suppressed the lethality of a subset of spindle assembly checkpoint gene knockouts, whereas Mps1 inhibitor treatment did not. Thus, despite Cdc20 ΔFL and Mps1 inhibitor-treated cells exhibiting similar effects on Spindle Assembly Checkpoint activity, as assessed by their mitotic arrest duration under anti-microtubule drug conditions ^7, 19^, they display distinct genetic dependencies, suggesting substantially differential contributions to the regulation of mitosis.

### The Cdc20 **Δ**FL cell line suppresses the lethality of Mad2 and other SAC protein knockouts

In addition to cases in which a gene knockout displayed an exacerbated growth defect in the Cdc20 ΔFL background, we also identified gene targets that displayed substantially improved growth in the Cdc20 ΔFL mutant cell line. These genes were strongly enriched for essential SAC genes. The Cdc20 ΔFL cell line suppressed the lethality of MAD2L1 (Mad2), TRIP13, and BUB1B (BubR1) knockouts (Fig. 1D). The most prominent change in growth behavior was seen for Mad2, with a CRISPR score of -3.40 in control HeLa cells, reflective of strong lethality, versus 0.088 in Cdc20 ΔFL cells. This enrichment for SAC genes was also reflected in K562 Cdc20 ΔFL cells, as these cells suppressed the lethality of a BUB1B knockout (Fig. S1G). In contrast, the addition of Mps1 inhibitor did not suppress the lethality of MAD2L1, TRIP13, or BUB1B gene knockouts (Fig. 2F, S1C).

To validate this unexpected growth behavior, we compared the effect of inducibly eliminating Mad2 in control HeLa cells or Cdc20 ΔFL cells on cellular fitness using the fluorescence-based competition assay. Although ΔMad2 cells were rapidly lost from the competition assay by Day 3, consistent with potent cellular lethality, Cdc20 ΔFL + ΔMad2 cells were able to effectively compete with Cdc20 ΔFL cells, with the ratio of Cdc20 ΔFL ΔMad2 vs. Cdc20 ΔFL cells plateauing at 0.7 (Fig. 3A). We also observed similar results using a lung adenocarcinoma-derived A549 cell line containing the Cdc20 ΔFL mutant (Fig. S3A). This growth behavior is consistent with Mad2 being dispensable for viability in the background of the Cdc20 ΔFL mutant. Indeed, in both the HeLa and A549 Cdc20 ΔFL cell line backgrounds, we were able to create stable clonal cell lines in which Mad2 was completely eliminated using CRISPR/Cas9-based gene targeting (Fig. 3B, S3B-D). Despite the complete suppression of lethality, the HeLa Cdc20 ΔFL + ΔMad2 cell line still lacked a functional spindle assembly checkpoint, as it was unable to maintain a mitotic arrest following treatment with the KIF11 inhibitor STLC (Fig. S3E), similar to Cdc20 ΔFL cells ^7^. This indicates that the loss of Mad2 does not compromise cellular viability in the Cdc20 ΔFL background despite the absence of a functional checkpoint. Using the cellular competition assay, we were also able to recapitulate the suppression of a BubR1 knockout in the HeLa Cdc20 ΔFL cell line, although not to the same extent as Mad2 (Fig. S3F). This partial suppression may reflect the additional roles of BubR1 in regulating chromosome segregation ^20–23^. Overall, these results demonstrate that a subset of checkpoint proteins become dispensable for viability in cells lacking the full length Cdc20 isoform.

**Fig. 3:**
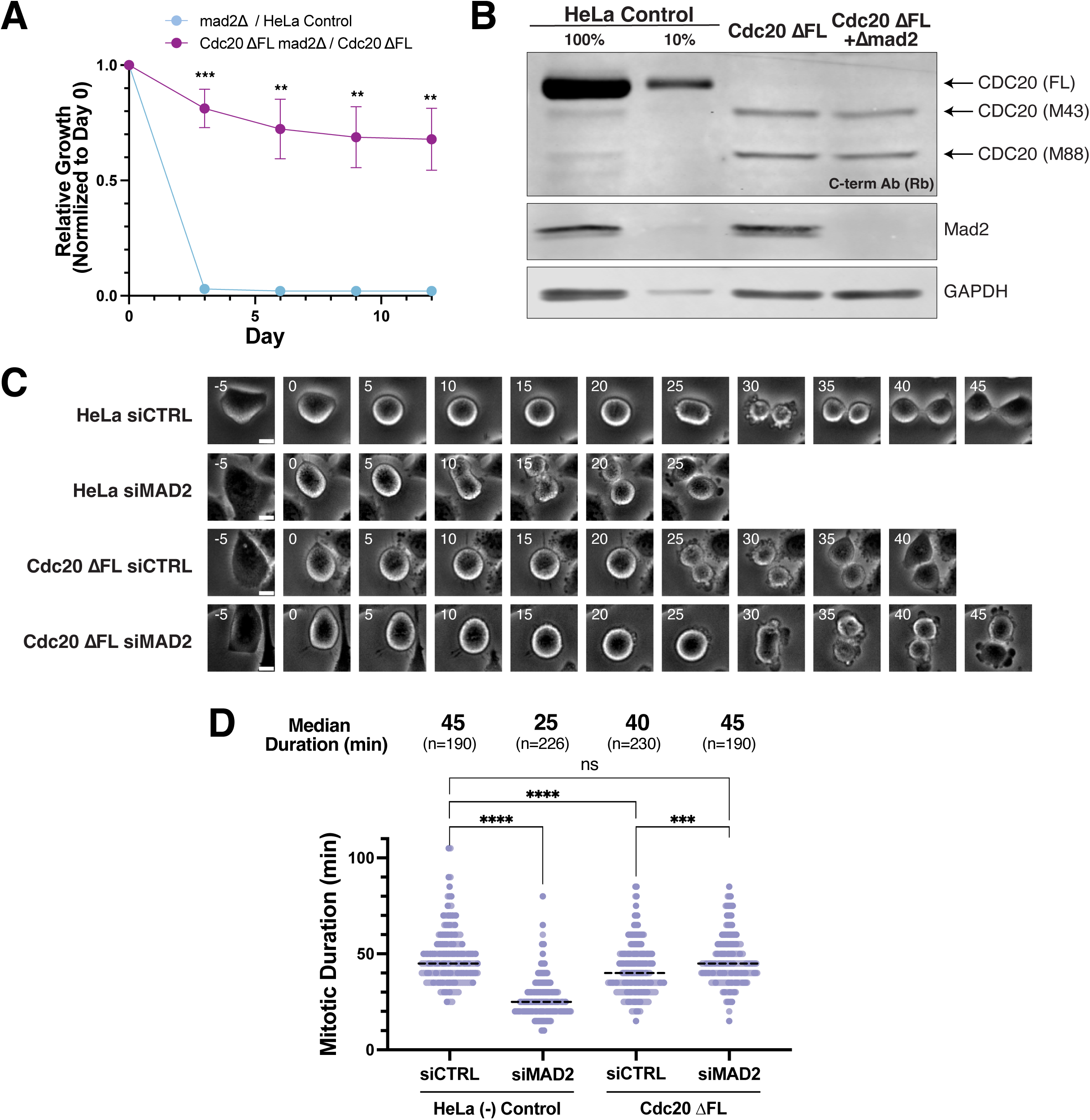
Mad2 is dispensable in Cdc20 ΔFL cells. **a,** Competitive growth assay to test the relative fitness of inducibly eliminating Mad2 in HeLa control or Cdc20 ΔFL cells (Clone #1). Data points reflect mean ± s.e.m; *n =* 3 experimental replicates. Student’s two-sample-*t-*tests with two-tailed distribution comparing relative ratios shown for Days 3-12. \*\**P*<0.01, \*\*\**P*<0.001. **b,** Western blot analysis of endogenous Cdc20 and Mad2 in HeLa, Cdc20 ΔFL (Clone #3), and Cdc20 ΔFL Δmad2 B3 cells (derived from the Cdc20 ΔFL Clone #3 cell line) using antibodies recognizing the C-terminus of human Cdc20 (aa 450-499). GAPDH was used as loading control. **c,** Representative stills of HeLa siCTRL, HeLa siMAD2, Cdc20 ΔFL siCTRL (Clone #1), and Cdc20 ΔFL siMAD2 (Clone #1) mitotic cells taken every 5 minutes from cell rounding at mitotic entry (t = 0 min) to cell flattening after mitotic exit. Scale bar = 10 µm. **d,** Quantification of mitotic timing (cell rounding at mitotic entry to cell flattening after mitotic exit) for HeLa and Cdc20 ΔFL (Clone #1) cells treated with siCTRL/siMAD2. Data are median across two experimental replicates; replicates are color coded. The total number of cells analyzed is indicated. The Kruskal-Wallis and Dunn’s post-hoc test was performed on the NEBD to onset of cyclin B1 degradation timings. \*\*\**P*<0.001, \*\*\*\**P*<0.0001, ns>0.9999.

### Mitotic progression is not delayed in Cdc20 **Δ**FL cells

Given the essential role of MAD2L1, the viability of Cdc20 ΔFL + ΔMad2 knockout cells was unexpected. We next sought to determine the basis for this suppression. We first considered whether this behavior could reflect partially compromised APC/C activity when full length Cdc20 is eliminated, resulting in a mitotic delay. Prior work found that impairing APC/C activity by depleting the E2 ubiquitin-conjugating enzymes UBE2S and UBE2C partially suppressed the lethality of Mad2 depletion ^24^. UBE2S + UBE2C-depleted cells display a delay in mitotic timing due to compromised APC/C activity ^24^, whereas Mad2-depleted cells undergo accelerated mitotic progression, entering anaphase prior to completely aligning their chromosomes. Thus, when these two perturbations are combined, mitotic duration is sufficiently delayed to allow adequate time for chromosome alignment even in the absence of Mad2 ^24^. In contrast, we found that Cdc20 ΔFL cells did not display an increased mitotic duration, with an overall timing of 40 minutes, similar to control cells (Fig. 3C,D). Furthermore, when Mad2 is then depleted by RNAi in Cdc20 ΔFL cells, we still observed a similar mitotic duration to control cells (45 minutes) (Fig. 3D). Therefore, the basis for the suppression of a Mad2 knockout lethality in the Cdc20 ΔFL cell line is not due to partially compromised APC/C activity counteracting the accelerated mitotic progression of a Mad2 depletion.

### Cyclin B1 degradation kinetics and activation timing are not strongly altered in Cdc20 **Δ**FL cells

To evaluate APC/C activity, we next measured the degradation of the APC/C substrate cyclin B1 by tagging CCNB1 with a Venus tag at its endogenous locus (Fig. 4A,B, S4A; ^25^). We first measured the kinetics of cyclin B1 degradation during anaphase, which provides a proxy for APC/C activity. Cells treated with the APC/C inhibitor proTAME displayed a notable decrease in the rate of cyclin B1 degradation (Fig. 4B). In contrast, both Cdc20 ΔFL and Cdc20 ΔFL + ΔMad2 cells displayed only modest changes in the rate of cyclin B1 degradation (Fig. 4B). Therefore, APC/C activity remains largely unaffected in Cdc20 ΔFL cells and cannot account for the robust suppression of the lethality from Mad2 depletion.

**Fig. 4:**
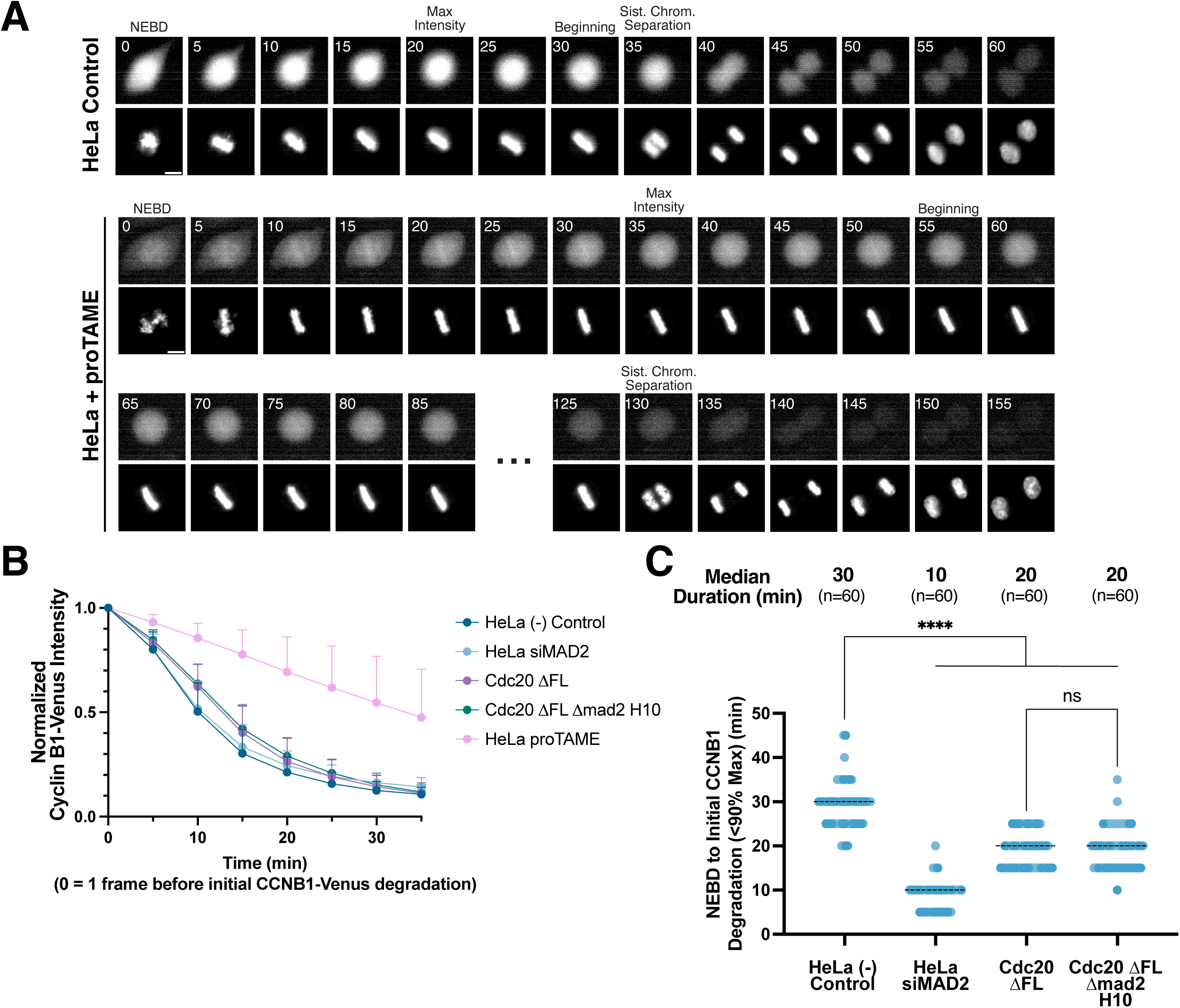
Cyclin B1 degradation kinetics and activation timing are not strongly altered in Cdc20 ΔFL cells. **a,** Representative stills of HeLa cells alone and HeLa cells treated with 12 µM proTAME from NEBD (t = 0 min) to 25 minutes after sister chromatid separation taken every 5 minutes. Images show Cyclin B1 endogenously tagged with Venus fluorescence and DNA (SIR-DNA). Max intensity is defined as the timepoint with maximum Cyclin B1-Venus fluorescence intensity. The onset of cyclin B1 degradation (Beginning) is defined here as when the Cyclin B1-Venus fluorescence intensity is <90% the maximum fluorescence intensity. **b,** Cyclin B1 degradation curves for HeLa, Cdc20 ΔFL (Clone #1), HeLa siMAD2, Cdc20 ΔFL Δmad2 H10 (derived from the Cdc20 ΔFL Clone #1 cell line), and HeLa proTAME (12 µM) cells during anaphase. Time t = 0 min is defined as 5 minutes before onset of cyclin B1 degradation (Beginning; see **a** for definition), and all time points (every 5 minutes for 35 minutes) are normalized to the Cyclin B1-Venus fluorescence intensity at t = 0. Data points reflect mean ± s.e.m.; *n* = ∼60 cells across two experimental replicates (exact number of cells seen in **c**). **c,** Quantification of timing from NEBD to onset of cyclin B1 degradation, as defined in **a**, for HeLa, Cdc20 ΔFL (Clone #1), HeLa siMAD2, Cdc20 ΔFL Δmad2 H10 cells. Data are median across two experimental replicates; replicates are color coded. The total number of cells analyzed is indicated. The Kruskal-Wallis and Dunn’s post-hoc test was performed on the NEBD to onset of cyclin B1 degradation timings. \*\*\*\**P*<0.0001, ns>0.9999.

To evaluate the regulation of APC/C activation, we next measured the timing of anaphase onset based on two criteria. First, we tested the time from Nuclear Envelope Breakdown (NEBD) to the onset of cyclin B1 degradation (Fig. 4A, B, S4A) as a proxy for the timing of APC/C activation. Second, we assessed the physical onset of anaphase based on the first timepoint at which sister chromatid separation could be observed. As expected, Mad2 depletion using 24 hours of RNAi treatment (see Methods) resulted in a drastic decrease in the time to initial cyclin B1 degradation of 10 minutes, compared to 30 minutes in control cells (Fig. 4C). Similarly, these Mad2-depleted cells exhibited an accelerated time from NEBD to sister chromatid separation, with timing of 15 minutes compared to 35 minutes in control cells (Fig. S4B). By comparison, Cdc20 ΔFL cells displayed only an intermediate initial cyclin B1 degradation timing (20 minutes) and sister chromatid separation timing (25 minutes) compared to control cells (Fig. 4C, S4B). Importantly, both initial cyclin B1 degradation timing (20 minutes, Fig. 4C) and sister chromatid separation timing (30 minutes, Fig. S4B) in Cdc20 ΔFL cells were unaffected by the further depletion of Mad2. This suggests that the Cdc20 ΔFL cells do not rely on Mad2 to regulate mitotic duration, and that there must be additional regulatory events that regulate the timing of APC/C activation.

### Cdk1-mediated phosphorylation of Cdc20 regulates the timing of APC/C activation

Prior work proposed that Mad2 acts as a mitotic timer by restraining the activity of Cdc20-bound APC/C until chromosomes are aligned ^5, 6, 26^. However, Cdc20 is also inhibited by Cyclin-dependent kinase (CDK)-dependent phosphorylation, which has been proposed to play roles in mitotic entry ^27, 28^ and the timing of anaphase events ^29,30^. The phosphorylation of Cdc20 prevents it from binding to and activating the APC/C ^29–31^. Thus, we next considered the contributions of Cdk1-mediated Cdc20 phosphorylation in regulating APC/C activation in Cdc20 ΔFL cells, which can no longer be regulated by the Mad2-containing MCC complex.

To test the contribution of Cdc20 phosphorylation to APC/C activation, we tested the consequences of a Cdc20 mutant in which four Cdk1 phosphorylation sites (S41, T55, T59, T70) were mutated to alanine to prevent their phosphorylation (Cdc20 4A; Fig. S5A). To conduct these studies, we used an RNAi-mediated strategy to replace wild type Cdc20 with this mutant ^7^. Based on the timing of initial cyclin B1 degradation, the Cdc20 4A mutant displayed only slightly accelerated anaphase onset with a timing of 20 minutes for NEBD to initial cyclin B1 degradation compared to 25 minutes in control replacement cells (Fig. 5A). The Cdc20 4A construct also displayed a decrease in the overall timing from NEBD to sister chromatid separation to 25 minutes compared to 35 minutes in control replacement cells (Fig. S5B) and displayed a modest increase in chromosome segregation defects (Fig. 5B, C, S5C, D). Importantly, APC/C activity remained unchanged based on the rate of cyclin B1 degradation during anaphase (Fig. S5E). Thus, in cells that contain full SAC activity, the phosphorylation state of Cdc20 makes a modest contribution to regulating the timing of APC/C activation.

**Fig. 5:**
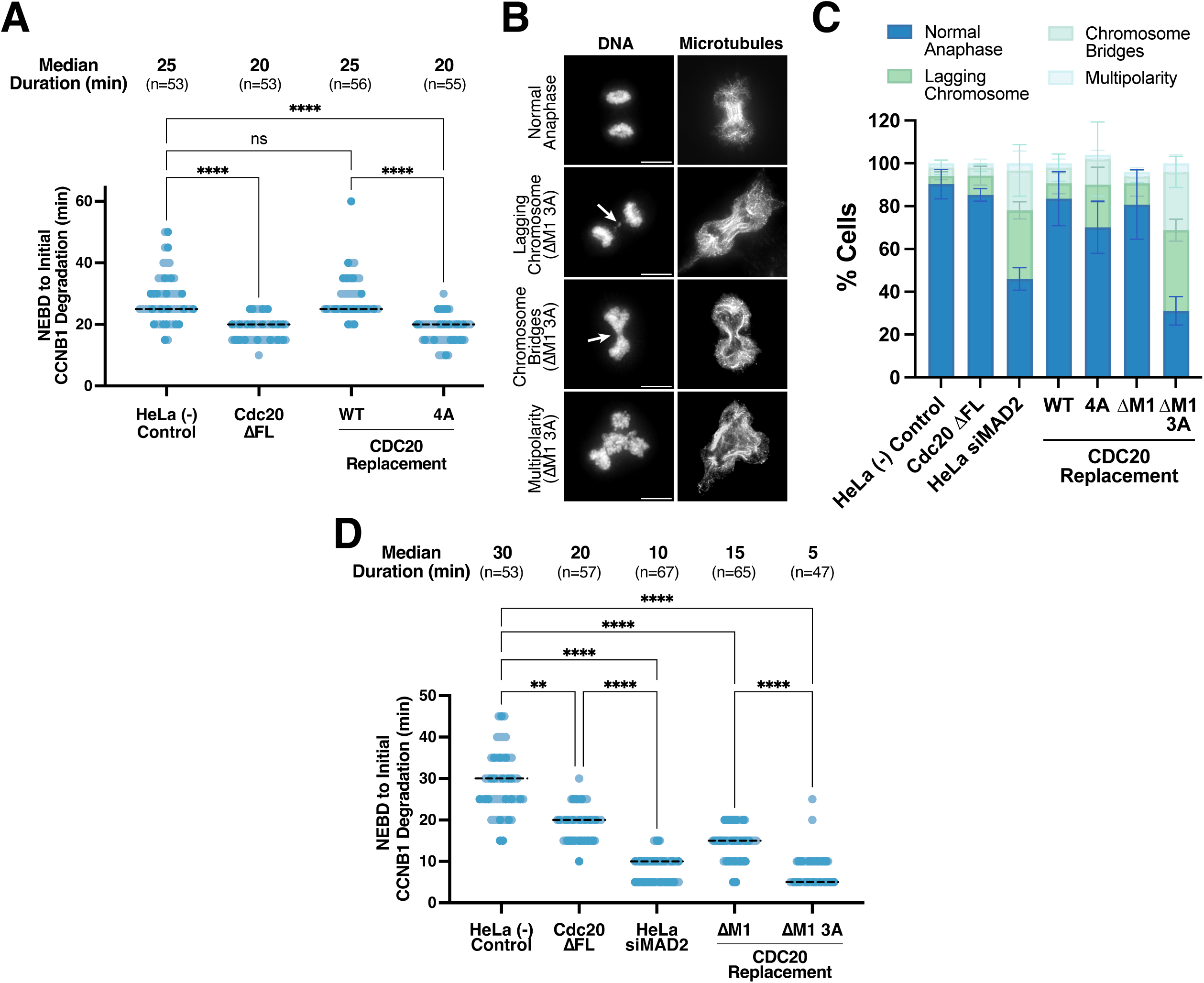
Cdk1-mediated phosphorylation of Cdc20 regulates the timing of APC/C activation. **a,** Quantification of timing from NEBD to onset of cyclin B1 degradation, as defined in **4a** for HeLa control cells, Cdc20 ΔFL (Clone #1) cells, and cells in which endogenous CDC20 protein is replaced with either WT siRNA-resistant *CDC20* cDNA (HeLa siCDC20 + TetON::CDC20 WT) or a mutant with four Cdk1-mediated phosphorylation sites (S41, T55, T59, T70) mutated to alanine (4A). Data are median across two experimental replicates; replicates are color coded. The total number of cells analyzed is indicated. **b,** Representative deconvolved Z-projected immunofluorescence images of anaphase cells from cell lines indicated in **c**. Images show microtubules (DM1α) and DNA (Hoeschst). Arrowheads indicate the specified anaphase error. Scale bars, 10 µM. **c,** Percentage of cells with anaphase defects in cells indicated. *n* > 150 cells per cell line, across three experimental replicates. Mean percentage is shown, error bars indicate SD. **d,** Quantification of timing from NEBD to onset of cyclin B1 degradation, as defined in **4a**, for HeLa control cells, Cdc20 ΔFL (Clone #1) cells, HeLa siMAD2 cells, and cells in which endogenous CDC20 protein is replaced with a mutant disrupting the Met1 start site (ΔM1) or a mutant disrupting the Met1 start site with also three Cdk1-mediated phosphorylation sites mutated (T55, T59, T70) to alanine (ΔM1 3A). Data are median across two experimental replicates; replicates are color coded. The total number of cells analyzed is indicated. For **a,d**, the Kruskal-Wallis and Dunn’s post-hoc test was performed on the NEBD to onset of cyclin B1 degradation timings. \*\**P*<0.01, \*\*\**P*<0.001, \*\*\*\**P*<0.0001.

### Coordinate control of mitotic timing by SAC and CDK activity

As our results suggest that SAC-dependent APC/C-Cdc20 inhibition and Cdk1-mediated phosphorylation of Cdc20 both act to delay the activation of the APC/C, we next tested the consequences of simultaneously removing both of these modes of regulation. For these experiments, we replaced endogenous Cdc20 with a ΔM1 3A construct, which lacks the full length Cdc20 isoform and has the three Cdk1-mediated phosphorylated sites mutated to alanine (3A; S41 is located upstream of the M43 start site) (Fig. S5F). Strikingly, we found that these cells completely lost their ability to regulate the timing of APC/C activation, with an NEBD to initial cyclin B1 degradation timing of only 5 minutes (Fig. 5D, S5G,H). This decrease in mitotic duration is consistent with the absence of a mitotic timer, with APC/C activation occurring just after mitotic entry. Consistent with this dramatic reduction in wait timing before initial cyclin B1 degradation, we also found that the Cdc20 ΔM1 3A cells displayed severe chromosome mis-segregation (Fig. 5C, S5D).

Thus, Cdc20 ΔM1 3A cells, which lack both the SAC-dependent inhibition of APC/C-Cdc20 (ΔM1) and the Cdk-dependent phosphorylation of Cdc20 (3A), display more severe phenotypes compared to cells lacking only one of these two modes of regulation (Fig. 5A, C-D). This suggests that the timing of anaphase onset is controlled by the combined activity of the SAC inhibition and Cdk1 phosphorylation to restrain the activity of Cdc20-bound APC/C, ensuring that the cell has sufficient time for chromosomes to align at the metaphase plate before the APC/C is activated and mitosis proceeds.

### The total levels of Cdc20 regulate the timing of APC/C activation

Finally, as the amount of active APC/C has important consequences for its ability to target substrates for degradation, we tested the impact of the levels of total Cdc20 in controlling the timing of mitotic progression. Previous studies have identified varying Cdc20 protein levels across cell lines and have found that partial Cdc20 depletion in several of these cell lines results in a prolonged metaphase duration ^32, 33^. Notably, Cdc20 ΔFL cells also display reduced levels of Cdc20 protein due to the absence of the full length Cdc20 isoform (Fig. 1A). Thus, to test the consequences of increasing Cdc20 levels in Cdc20 ΔFL cells, we ectopically expressed constructs containing Cdc20 (Fig. S6A). Increasing the levels of all Cdc20 isoforms (including the checkpoint sensitive full length Cdc20 isoform containing the Box 1 motif) in the Cdc20 ΔFL cell line did not alter the wait timing nor overall anaphase onset (Fig. 6A, S6B). In contrast, overexpression of only the M43 and M88 Cdc20 isoforms in the Cdc20 ΔFL cell line (Fig. S6A) accelerated the timing from NEBD to initial cyclin B1 degradation from 20 minutes to 10 minutes (Fig. 6A) and the timing from NEBD to sister chromatid separation from 25 minutes to 15 minutes (Fig. S6B). Thus, when the full length Cdc20 isoform is present, Cdc20-Mad2 interactions likely buffer the effects of increased levels of Cdc20. However, when the SAC-deficient Cdc20 isoforms are overexpressed, this results in premature cyclin B degradation due to the absence of Mad2 interactions. Indeed, if we once again removed the regulation of the SAC by depleting Mad2 from cells overexpressing all of the Cdc20 isoforms, the initial cyclin B1 degradation and sister chromatid separation timings accelerated back to 10 minutes and 15 minutes, respectively (Fig. 6A, S6B).

**Fig. 6:**
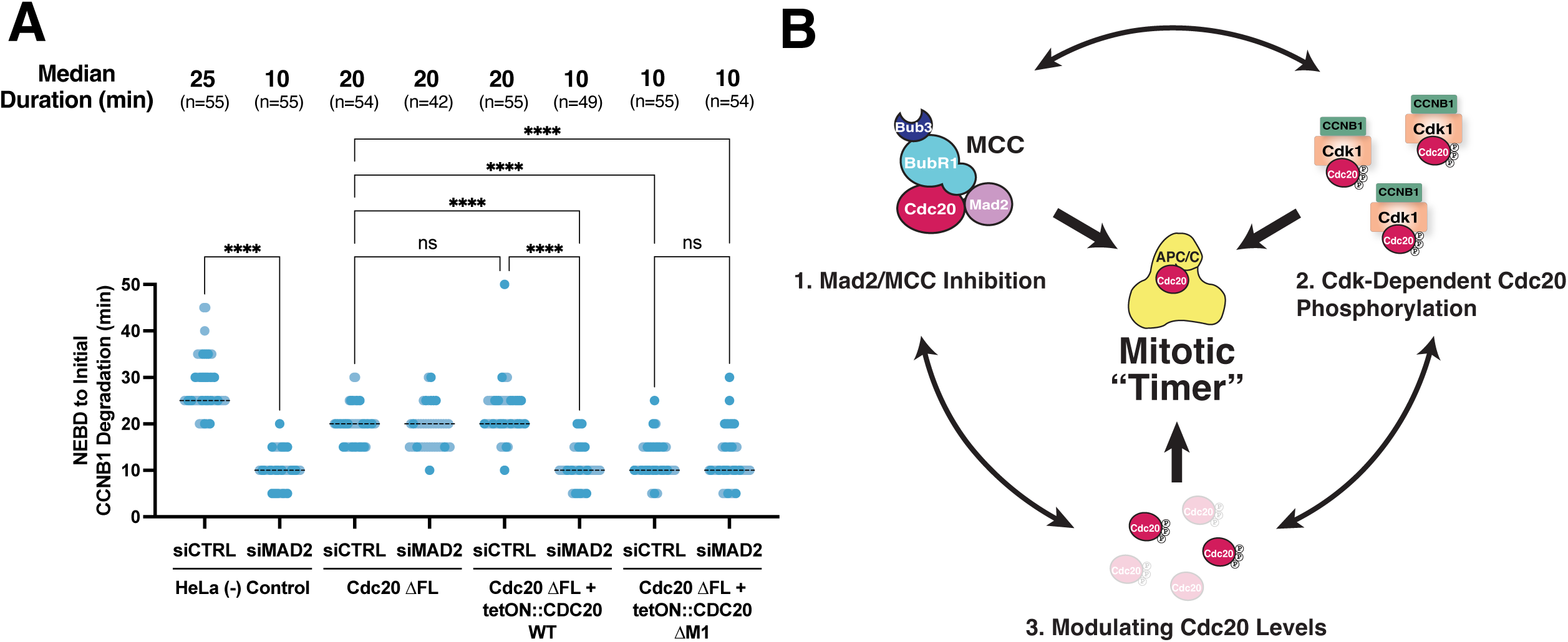
Cdc20 protein levels modulate a feedback loop involving Cdc20 and CDK activity that regulates the timing of APC/C activation. **a,** Quantification of timing from NEBD to onset of cyclin B1 degradation, as defined in **4a**, for HeLa control cells, Cdc20 ΔFL (Clone #1) cells, HeLa siMAD2 cells, and cells treated with 50 ng/mL doxycycline to induce expression of indicated *CDC20* constructs and RNAi against Mad2 or a non-targeting control pool. Data are median across two experimental replicates; replicates are color coded. The total number of cells is indicated. The Kruskal-Wallis and Dunn’s post-hoc test was performed on the NEBD to onset of cyclin B1 degradation timings. \*\*\**P*<0.001, \*\*\*\**P*<0.0001, ns>0.9999. **b,** Schematic of the model for factors involved in defining the “mitotic timer”.

Together, our results suggest that the timing of anaphase onset is controlled by three factors: 1) the presence of Mad2 and its inhibition of Cdc20, 2) Cdc20 inhibition by CDK-dependent phosphorylation, and 3) Cdc20 protein levels (Fig. 6B). Changes in any of these factors across physiological conditions would be expected to alter mitotic progression, with changes to a given pathway buffered by the presence of the others.

## Discussion

Previous studies have proposed multiple functions for Spindle Assembly Checkpoint proteins. In addition to the classically defined function of the checkpoint as a surveillance mechanism that is only activated in the presence of errors, the Spindle Assembly Checkpoint has been proposed to act as a regulator of chromosome segregation fidelity to control kinetochore-microtubule attachments, and as a mitotic timer that restricts mitotic progression to ensure that chromosomes have sufficient time to align at the metaphase plate. Here, we utilized a SAC-deficient Cdc20 ΔFL cell line that lacks the longer translational protein isoform of Cdc20 as a unique genetic background to explore the functional contributions of the SAC in human cells. Using CRISPR/Cas9-based functional genetics, we demonstrate that Cdc20 ΔFL cells display a synthetic lethal relationship with gene targets whose loss results in increased chromosome segregation defects, such as CENPE and MKI67. These genetic interactions are consistent with the classically defined role for the Spindle Assembly Checkpoint as a surveillance pathway that is only needed in the presence of mitotic errors, similar to work in budding yeast.

Strikingly, despite its essential role in control cells, we demonstrate that Mad2 becomes dispensable for viability in the Cdc20 ΔFL background. Our results indicate that Cdc20 ΔFL cells exhibit mostly unchanged APC/C activity and are still able to regulate the timing of APC/C activation, even when Mad2 is depleted from the cells. Therefore, although Mad2 is an important factor for the “mitotic timer”, our results in the Cdc20 ΔFL cells indicate that additional mechanisms also act to regulate mitotic progression. Specifically, we find that Cdk1-mediated phosphorylation of Cdc20 and the overall levels of Cdc20 are important in regulating APC/C activation. Cdk1-mediated phosphorylation regulates APC/C activation by preventing Cdc20 from activating the APC/C. However, the APC/C also regulates CDK activity by promoting the degradation of cyclin B, thereby inactivating CDK. This reciprocal inhibition would create a feedback loop in which partial Cdc20 activation and cyclin B destruction would result in reduced Cdc20 phosphorylation and the further activation of the APC/C-Cdc20 (Fig. S6C; see also ^30, 34^). This positive feedback loop for APC/C activation would depend at least partially on the levels of Cdc20 and the relative level of CDK activity. Therefore, this could explain in part why the total levels of Cdc20 protein tune the initiation of APC/C activation and regulate the timing of anaphase onset (Fig. 6A). Decreasing overall Cdc20 levels would decrease the availability of unphosphorylated Cdc20, delaying the partial activation of the APC/C and the initiation of the positive feedback loop. Consistent with this model, we found that CRISPR-Cas9 mediated knockouts of the mitotic cyclins CCNB1 or CCNB2, which would reduce CDK activity, are synthetic lethal in Cdc20 ΔFL cells (Fig. 1D).

In summary, our work demonstrates that the precise timing of anaphase onset depends on the interdependent and overlapping actions of: (1) Mad2 inhibition of Cdc20-bound APC/C activity, (2) Cdk1-mediated phosphorylation and inhibition of Cdc20, and (3) Cdc20 protein levels modulating the initiation of a feedback loop involving Cdc20 and CDK activity. These three factors work together in regulating APC/C activation, with changes to one given pathway buffered by the actions of the other pathways. Simultaneously perturbing these pathways, as seen in Cdc20 ΔM1 3A cells that lack both the SAC-dependent inhibition of APC/C-Cdc20 and the CDK-dependent phosphorylation of Cdc20, results in near immediate mitotic exit and catastrophic chromosome mis-segregation (Fig. 5A, C-D), highlighting the importance of all three factors in regulating APC/C activation. This expanded understanding of the control of mitotic is important for predicting tumor sensitivity in cases where the Spindle Assembly Checkpoint is weakened. For example, Mps1 inhibitors are currently in clinical trials for the treatment of solid cancers (NCT02792465, NCT03568422, NCT05251714) and recent work found that Cdc20 expression is a major predictor for Mps1 inhibitor sensitivity in aneuploid cancer cells ^33^, thereby implicating the Cdc20 levels as an important metric for guiding treatment. By analyzing cells lacking the full length Cdc20 translational isoform and SAC activity, our work expands the understanding of the regulation of mitotic timing.

## Materials & Methods

### Tissue Culture and Reagents

HeLa, A549, and HEK293T cells were cultured in Dubelcco’s modified Eagle medium (DMEM) supplemented with 10% fetal bovine serum (FBS), 100 U/mL penicillin and streptomycin, and 2 mM L-glutamine at 37°C under 5% CO_2_. All doxycycline-inducible cell lines were cultured in medium containing FBS certified tetracycline-free and were induced by the addition of 1 µg/mL doxycycline hyclate (Sigma-Aldrich) for 2 days. Cells were regularly monitored for mycoplasma contamination using commercial detection kits.

K562 cells were cultured in RPMI-1640 medium supplemented with 10% FBS and 100 U/mL penicillin and streptomycin at 37°C under 5% CO_2_. K562 cells were regularly monitored for mycoplasma contamination using commercial detection kits.

Other drugs used on human cells were STLC (Sigma Aldrich; 10 µM), proTAME (APC/Ci; R&D Systems, 12 µM), and AZ-3146 (Mps1i; Selleck Chemicals, 0.5-1.75 µM). The antibodies used in this study are described in the relevant methods and in Supplementary Table 2.

### Virus Production, Transduction, and Selection

To generate lentivirus, HEK293T cells were seeded at ∼75% confluency in a 6-well plate and then cultured for 16 hours at 37°C. 1.2 µg lentiCas9-sgRNA transfer plasmid, 1 µg psPAX2 packaging plasmid, and 0.4 µg vsFULL envelope plasmid using Xtremegene-9 (Roche) were then transfected into these cells. After 16 hours, fresh medium was replaced. After 24 hours, the now virus-containing medium was collected and stored at - 80°C.

For lentiviral transduction, cells of interest were seeded at ∼25% confluency (Competition assays) or 12.5% confluency (Mad2 knockout cell line production) and cultured for 16 hours at 37°C. Fresh medium supplemented with 20 µg/mL polybrene was replaced, and 75-100 µL of lentivirus was then added to the cells. Cells then underwent a spinfection (2250 RPM, 45 mins, 37°C), then were cultured for 16 hours. Fresh medium was then applied, and cells were then further cultured for 8 hours. Cells were then passaged and selected with 0.5 µg/mL puromycin for 64-72 hours.

The following RNA sequences were used: sgHS1: 5’-GCCGATGGTGAAGTGGTAAG-3’ sgMAD2L1: 5’-GTATTTCTGCACTCGAGTAA-3’, sgCENPE: 5’-TCAGCCTGATAGGATGGCGG-3’. sgMKI67: 5’-ACAGAGGTTCCTAAGAGAGG-3’, sgBUB1B: 5’-GAGCAGAACTATCCTCAAGG-3’.

### Cdc20 ***Δ***FL Cell Line Production

The Cdc20 ΔFL (Clone #1) cell line was first generated in ^7^. An extended protocol for how this cell line was generated can be found in the previous work. Briefly, for the Cdc20 ΔFL (Clone #1) cell line, HeLa cells were transfected with a pX330-based plasmid ^35^ expressing both spCas9 and the sgExon1 guide RNA (5’-CCTGCACTCGCTGCTTCAGC-3’) using X-tremeGENE-9 (Roche). A mCherry-expressing plasmid was also transfected in. Single mCherry-positive cells were sorted into 96-well plates using a BD FACSAria III sorter (BD Biosciences). Clones were then screened for successful gene editing (a premature stop codon introduced after Met1 but before Met43), and the nucleotide sequence of the *CDC20* alleles was determined by next-generation sequencing (Genewiz Amplicon-EZ).

The Cdc20 ΔFL (Clones #2,3) cell lines were generated in this study. HeLa cells were transfected with 1.25 µg pX330-based plasmid ^35^ expressing spCas9, the sgExon1 guide RNA (5’-CCTGCACTCGCTGCTTCAGC-3’), and BFP, and 1.25 µg pCR4-TOPO-based recombination plasmid containing *CDC20* sequence with a 2 nt nonsense mutation at the L13 residue and a 1 nt mutation at the sgExon1 guide PAM site using Lipofectamine 2000 (ThermoFisher). Homology arms of ∼500 bp on either side was used in the recombination template. Fresh medium was replaced 16 hours post transfection, then cells were allowed to row for 72 hours. Single BFP-positive cells were sorted into 96-well plates using a BD FACSAria III sorter (BD Biosciences). Clones were then screened for successful gene editing by first selecting for clones that failed to mitotically arrest after 16 hours of 10 µM STLC treatment. Clones were then screened for those lacking full length Cdc20 by Western blot analysis. The nucleotide sequence of the *CDC20* alleles was then determined by Sanger sequencing (Azenta).

K562 Cdc20 ΔFL and A549 Cdc20 ΔFL cell lines were generated in this study. K562 or A549 cells were transfected with a pX330-based plasmid ^35^ expressing both spCas9 and a sgExon1 guide RNA (K562: 5’ – GCGCTGGCAGCGCAAAGCCA – 3’, A549: 5’-CCTGCACTCGCTGCTTCAGC-3’) using X-tremeGENE-9 (Roche). A mCherry-expressing plasmid was also transfected in. Single mCherry-positive cells were sorted into 96-well plates using a BD FACSAria III sorter (BD Biosciences). Clones were then screened for successful gene editing by first selecting for clones that failed to mitotically arrest after 16 hours of 10 µM STLC treatment. The nucleotide sequence of the *CDC20* alleles was then determined by Sanger sequencing (Azenta).

### Mad2 Knockout Cell Line Production

Mad2 knockout stable clonal cell lines were obtained by lentiviral transduction of a sgRNA targeting the *MAD2L1* gene (sgMAD2L1, 5’-GTATTTCTGCACTCGAGTAA-3’) cloned into the pLCV2-Opti plasmid ^36^. Cells were selected with 0.5 µg/mL puromycin (Gibco) for 64-72 hours. Single cells were sorted into 96-well plates using flow cytometry. Clones were screened for successful knockout by Western Blot and confirmed by sequencing the nucleotide sequence of the *MAD2L1* allele using Sanger sequencing (Azenta).

### Production of Endogenously Venus-Tagged Cyclin B1 Cell Lines

Cell lines of interest were transfected with 1.25 µg of two pX330-based BFP+ plasmids expressing spCas9 and single-guide RNAs (gRNA sequences: 5’-ACTAGTTCAAGATTTAGCCA-3’, 5’-TGTAACTTGTAAACTTGAGT-3’) and 2.5 µg of a pCR4-TOPO CCNB1 (Exon 9)-GGSGSGGGSG Linker-Venus recombination plasmid using Lipofectamine 2000 (ThermoFisher). Homology arms of ∼500 bp on either side was used in the recombination template. Fresh medium was swapped 16 hours post transfection, then cells were allowed to grow for 72 hours. Venus positive cells were bulk sorted using a BD FACSAria III sorter (BD Biosciences).

### Production and Expression of Doxycycline-Inducible TetON:CDC20 Overexpression or Gene Replacement Cell Lines

Doxycycline-inducible cell lines were generated by homology-directed insertion into the *AAVS1* “safe-harbor” locus. 500 ng of a donor plasmid containing a hygromycin selection marker, tetracycline-responsive promoter, the *CDC20* transgene of interest, and reverse-tetracycline-controlled transactivator, with *AAVS1* homology arms on either side ^37^, and 500 ng of a pX330-based plasmid ^35^ expressing *sp*Cas9 and a guide RNA specific for the *AAVS1* locus (5’-GGGGCCACTAGGGACAGGAT-3’) were transfected into either HeLa or Cdc20 ΔFL (Clone #1) cells, using Lipofectamine 2000 (ThermoFisher). Fresh medium was replaced 16 horus post transduction, then allowed to grow for 24 hours. Cells were then selected with hygromycin (375 µg/mL) for 10-14 days.

For *TetON::CDC20* overexpression cell lines, cells were induced with 1 µg/mL doxycycline hyclate (Sigma-Aldrich) at 24 hour intervals for 2 days. For gene replacement cell lines, siCDC20 was forward transfected at cell seeding, with the transfection medium also containing 1 µg/mL doxycycline hyclate to express the ectopic inducible *CDC20* construct. Cells were induced for 2 days, with media containing 1 µg/mL doxycycline hyclate replaced at 24 hour intervals.

All of the plasmids and cell lines used in this study are listed in Supplementary Table 3 and 4, respectively.

### RNAi treatment

Custom siRNAs against *CDC20* (5′-CGGAAGACCUGCCGUUACAUU), *MAD2* (5′-UACGGACUCACCUUGCUUGUU), and a non-targeting control pool (D-001206-13) from Dharmacon were used in this study. siRNAs were applied at a final concentration of 50 nM. Exceptions are indicated in the figure or figure legend. A total of 2.5 µL Lipofectamine RNAiMAX (Invitrogen) was used per mL of the final transfection medium. For all experiments, siRNA treatment against *CDC20* was conducted for 48 hours to fully deplete endogenous Cdc20, and against *MAD2* and the non-targeting control pool for 24 hours, such that most endogenous Mad2 had been depleted but before the accumulation of defects caused by the depletion would lead to widespread cell death.

siRNAs were transfected during cell seeding except for additional partial Mad2-depletion conducted in doxycycline-inducible *CDC20* overexpression experiments. In those experiments, RNAi against *MAD2* was reverse transfected 24 hours after cell seeding. The medium was swapped 8 hours later, and cells were cultured for an additional 16 hours.

### Generation of Lentiviral sgRNA Libraries and Large-Scale Lentiviral Production

We used a generated lentiviral sgRNA library described in ^16^ comprising 14,989 unique sgRNA sequences targeting 1411 genes. See ^16, 38^ for an extended protocol on how the library was generated and for large-scale lentiviral production.

### CRISPR Screen

For the screening of HeLa and Cdc20 ΔFL (Clone #1) cells, lentivirus containing the targeted lentiviral sgRNA library was transduced into 39 million cells. Cells were then cultured for 2 days before passaging into DMEM medium containing 0.4 µg/mL puromycin and were then selected for 4 days. At the end of selection, pellets of 5 million cells were taken to quantify initial sgRNA library representation. Cells were then passaged every 2 days for 14 population doublings. For the Mps1 inhibitor treatment condition, cells were grown in a low dose (0.5-1.75 µM) of the AZ-3146 Mps1 inhibitor (Selleck Chemicals) that would still impede SAC function but would only reduce overall cell proliferation by ∼5%, allowing us to identify genes whose knockout effects on cellular viability are either exacerbated or suppressed due to weakened Mps1 activity. Pellets of 5 million cells were collected at the 14 population doublings to quantify final sgRNA library representation.

For the screening of K562 and K562 Cdc20 ΔFL cells, lentivirus containing the targeted lentiviral sgRNA library was transduced into 35 million cells. Cells were then cultured for 2 days before passaging into RPMI-1640 medium containing 3 µg/mL puromycin and selection for 6 days. At the end of selection, pellets of 5 million cells were taken to quantify initial sgRNA library representation. Cells were then passaged every 2 days for 14 population doublings. Pellets of 5 million cells were collected at 14 population doublings to quantify final sgRNA library representation.

### CRISPR Screen Sequencing and Analysis

Please see ^16^ for an extended protocol on how the sequencing library was prepared from cells collected during the CRISPR screen. Sequencing was done for 50 cycles on an Illumina Hiseq 2500 using the following primers:

Read 1 sequencing primer (secondary): 5′-GTTGATAACGGACTAGCCTTATTTAAACTTGCTATGCTGTTTCCAGCATAGCTCTTAAAC-3′

Index sequencing primer: 5′-TTTCAAGTTACGGTAAGCATATGATAGTCCATTTTAAAACATAATTTTAAAACTGCAAACTACCCAAGAAA-3′

High-throughput sequencing reads from the CRISPR screens were then mapped to the sgRNA library using Bowtie. MAGeCK-RRA ^39^ was then used to generate gene scores representing the median log_2_ fold change in final sgRNA abundance over initial sgRNA abundance.

For comparisons between cell lines, the differential scores were calculated using the median log_2_ fold changes of the condition of interest and the (-) control condition.

### Western Blotting

Cells were washed in PBS and pelleted before frozen at -80° C until lysis. Cell pellet was lysed on ice for 25 minutes in fresh urea lysis buffer (50 mM Tris pH 7.5, 150 mM NaCl, 0.5% NP-40, 0.1% SDS, 6.5 M Urea, 1 x Complete EDTA-free protease inhibitor cocktail (Roche), 1 mM PMSF). Lysates were centrifugated and supernatant was collected and mixed with Laemmli sample buffer. Protein concentrations were measured using Pierce^TM^ BCA Protein Assay Kit (Thermo Fisher). Lysates were heated at 95° C for 5 minutes, and 20-30 µg of total protein was separated by SDS-PAGE on a 10% or 12.5% acrylamide gel and was then transferred to PVDF membrane (Cytiva).

Membrane was blocked for 30 minutes in blocking buffer (2% milk in PBS + 0.1% Tween-20). Primary antibodies were diluted in blocking buffer and applied to the membrane overnight at 4° C. Membrane was then washed with PBS + 0.1% Tween-20. IRDye 800CW goat anti-mouse, IRDye 680RD goat anti-mouse, IRDye 800CW goat anti-rabbit, and/or IRDye 680RD goat anti-rabbit secondary antibodies, depending on which primary antibodies were used, were diluted in blocking buffer and applied to the membrane for 1 hour at room temperature. Membrane was then washed with PBS + 0.1% Tween-20 and then subsequently PBS. Membrane was then imaged using the Odyssey CLx Imager (LI-COR) and subsequently analyzed using LI-COR Image Studio software. For Western blot analysis of K562 cells, standard chemiluminescence was conducted instead of imaging by LI-COR. For standard chemiluminescence, instead of applying IRDye secondary antibodies, HRP-conjugated secondary antibodies (GE Healthcare or Kindle Biosciences) were diluted in 0.2% milk in PBS + 0.05% Tween-20 and applied to the membrane for 1 hour at room temperature. After washing in PBS + 0.05% Tween-20, clarity-enhanced chemiluminescence substrate (Bio-Rad) was added to the membrane according to the manufacturer’s instructions. Membranes were imaged using the KwikQuant Imager (KindleBiosciences; Exposure time 15 seconds) and analyzed using Adobe Photoshop.

The following primary antibodies were used: anti-C terminus (amino acids 450-499) of human Cdc20 (C-term Ab, 1:500, Abcam, ab26483), anti-N terminus (amino acids 1-175) of human Cdc20 (N-term Ab, 1:300, Santa Cruz Biotechnology, sc-13162), anti-Mad2 (1:5000, Abcam, ab97777), anti-GAPDH (1:1000, Santa Cruz Biotechnology, sc-47724), anti-alpha tubulin (1:5000, Sigma-Aldrich, T9026), β-actin (1:20,000, Santa Cruz Biotechnology, sc-47778, HRP-conjugated). The following secondary antibodies were used: IRDye 680RD Goat anti-Rabbit (LI-COR 92668071), IRDye 680RD Goat anti-Mouse (LI-COR 92668070), IRDye 800CW Goat anti-Rabbit (LI-COR 92632211), IRDye 800CW Goat anti-Mouse (LI-COR 92632210) all at 1:10,000 dilution in 2% milk in PBS + 0.1% Tween-20.

The uncropped original images of all western blots in this study are provided in Supplementary Figure 1.

### Immunofluorescence microscopy

Cells were seeded in 12-well on poly-L-lysine-coated (Sigma-Aldrich) coverslips and cultured for 24 hours prior to fixation. Cells were fixed in 4% formaldehyde in PHEM (60 mM PIPES, 25 mM HEPES, 10 mM EGTA, and 4 mM MgSO_4_) for 10 min at 37°C followed by three washes in PBS + 0.1% Triton X-100. Cells were then blocked in AbDil (20 mM Tris-HCl pH 7.5, 150 mM NaCl, 0.1% Triton X-100, 3% BSA) for 30 min at room temperature or overnight at 4°C. Primary antibodies were diluted in AbDil and applied to the coverslips for 1 hour at room temperature. Cells were then washed three times in PBS + 0.1% Triton X-100. Secondary antibodies were diluted 1:300 in AbDil with 0.3 µg/mL Hoeschst-33342 (Invitrogen) and applied to the coverslips for 45 min at room temperature. Cells were washed three times in PBS + 0.1% Triton X-100, then mounted in p-phenylamine diamine (PPDM) onto coverslips. Immunofluorescence cell images were acquired on a DeltaVision Core deconvolution microscope (Applied Precision) equipped with a CoolSnap HQ2 CCD camera. Approximately 30 Z-sections were acquired at 0.2 µm steps using a 60x, 1.42 NA Olympus U-PlanApo objective. For representative images, Z-projected immunofluorescence images were deconvolved using the Softworx software.

The following primary antibodies were used: DM1α (1:1000, Cell Signaling, 3873). The following secondary antibodies were used: Donkey Anti-Mouse IgG (H+L) AlexaFluor488 (1:300, Invitrogen, A21206).

### Time Lapse Cell Rounding Experiments for Mitotic Timing

Cells were seeded in 12-well polymer-bottomed plates (Cellvis, P12-1.5P) at 12.5% confluency and then cultured for 24 hours. For cells depleted by RNAi, RNAi treatment was conducted at the time of cell seeding. Media was then replaced with CO_2_-independent media (Gibco) supplemented with 10% FBS, 100 U/mL penicillin and streptomycin, and 2 mM L-glutamine. For mitotic arrest timing analysis, media was additionally supplemented with 10 µM STLC and cells were allowed to incubate for 1 hour before beginning time lapse. Images were acquired using a Nikon eclipse microscope equipped with a sCMOS camera (ORCA-Fusion BT, Hamamatsu) with a Plan Fluor 20x/0.5 NA objective at 5-minute intervals. Time lapse videos were obtained by acquiring phase-contrast images for 20 hours in 5-minute intervals (48 hours for mitotic arrest timing analysis). The image brightness and contrast were first adjusted in ImageJ. Pixel-based classification was then performed on each adjust image to create probability map images. These were then processed using CellProfiler ^40^ to identify and track mitotic cells. Mitotic duration was determined as the time from cell rounding at mitotic entry to cell flattening after mitotic exit; durations were confirmed manually. For mitotic arrest timing analysis, only cells that enter mitosis in the first 24 hours were analyzed.

### Time Lapse Cyclin B1 Degradation

Cells were seeded in 12-well polymer-bottomed plates (Cellvis, P12-1.5P) such that at beginning of time lapse, cells were at 25% confluency. Media was then replaced with CO_2_-independent media (Gibco) supplemented with 10% FBS, 100 U/mL penicillin and streptomycin, 2 mM L-glutamine, and 1 µM SiR-DNA (Cytoskeleton Inc.), followed by a 1 hr incubation at 37°C. Images (Yellow Fluorescent Protein [YFP] and Cy5) were acquired at 5-minute intervals for 24 hours using the appropriate filter settings on a Nikon eclipse microscope equipped with a sCMOS camera (ORCA-Fusion BT, Hamamatsu) with a Plan Fluor 20x/0.5 NA objective.

Cells undergoing mitosis were selected in Fiji (ImageJ, NIH) and then analyzed using a custom CellProfiler pipeline ^40^. The pipeline segmented cells based on SiR-DNA signal, performed background subtraction, and measured nuclear Cyclin B1-Venus fluorescence intensity in each cell. The onset of cyclin B1 degradation (Beginning) was defined as the timepoint with a decrease in CCNB1-Venus fluorescence intensity from the previous timepoint > 10%. One exception was for experiments that included cells treated with 12 µM proTAME, as a decrease in CCNB1-Venus fluorescence intensity from the previous timepoint > 10% was not an accurate approximation for the onset of cyclin B1 degradation. For these experiments, the onset of cyclin B1 degradation (beginning) was defined instead as the timepoint in which the CCNB1-Venus fluorescence intensity is <90% the maximum fluorescence intensity of a given cell. The timing from Nuclear Envelope Breakdown (NEBD; as determined by DNA condensation) to onset of cyclin B1 degradation and the timing from NEBD to sister chromatid separation were quantified. To determine the rate of cyclin B1 degradation kinetics, time t = 0 min was set to one frame before onset of cyclin B1 degradation, and all time points were then normalized to the CCNB1-Venus fluorescence intensity at t = 0 min. The cyclin B1 degradation curves depict cells from time t = 0 min to five frames after sister chromatid separation.

### Competition Assay

To create BFP- and mCherry-expressing HeLa, Cdc20 ΔFL (Clone #1), A549, and A549 Cdc20 ΔFL cells, cells were infected with either eBFP2 lentivirus or mCherry lentivirus. Medium was replaced 16 hours post infection, then cells were allowed to grow for 72 more hours. BFP- and mCherry-positive cells were bulk sorted using a BD FACSAria III sorter (BD Biosciences).

Single-guide RNAs targeting either the control *LBR* gene (sgHS1) at a single site ^41^ or a gene of interest (sgGeneX) were cloned into the pLCV2-opti plasmid and introduced into BFP- and mCherry-expressing HeLa, Cdc20 ΔFL (Clone #1), A549, or A549 Cdc20 ΔFL cells by lentiviral transduction. BFP- and mCherry-expressing cells were then mixed 1:1 in pairwise assays. Proliferation of BFP- and mCherry-expressing cells was measured by flow cytometry on an LSR Fortessa (BD Biosciences) flow cytometer and analyzed using FlowJo every 3 days for 9 or 12 days. Cells were passaged when measurements were taken. The relative ratio of cells expressing pLCV2-opti-sgGeneX over cells expressing pLCV2-opti-sgHS1 was quantified and normalized first to the initial ratio when cells were first mixed (day 0), and then to normalized relative ratio of control cells (BFP- and mCherry-positive cells both expressing pLCV2-opti-sgHS1) at the same timepoint.

The following sgRNAs were used in competition assays: sgHS1: 5’-GCCGATGGTGAAGTGGTAAG-3’, sgMAD2L1: 5’-GTATTTCTGCACTCGAGTAA-3’, sgCENPE: 5’-sgCENPE: 5’-TCAGCCTGATAGGATGGCGG-3’. sgMKI67: 5’-ACAGAGGTTCCTAAGAGAGG-3’, sgBUB1B: 5’-GAGCAGAACTATCCTCAAGG-3’.

### Quantification and Statistical Analysis

Quantification of fluorescence intensity was done on unprocessed, maximally projected images using FIJI (Image J, NIH). Images for each experiment were acquired using the same microscope and acquisition settings for comparison. Representative images Statistical analyses were performed using Prism (GraphPad Software), with details of statistical tests and samples sizes for each experiment provided in the corresponding figure legend.

---

**Fig. S1:**
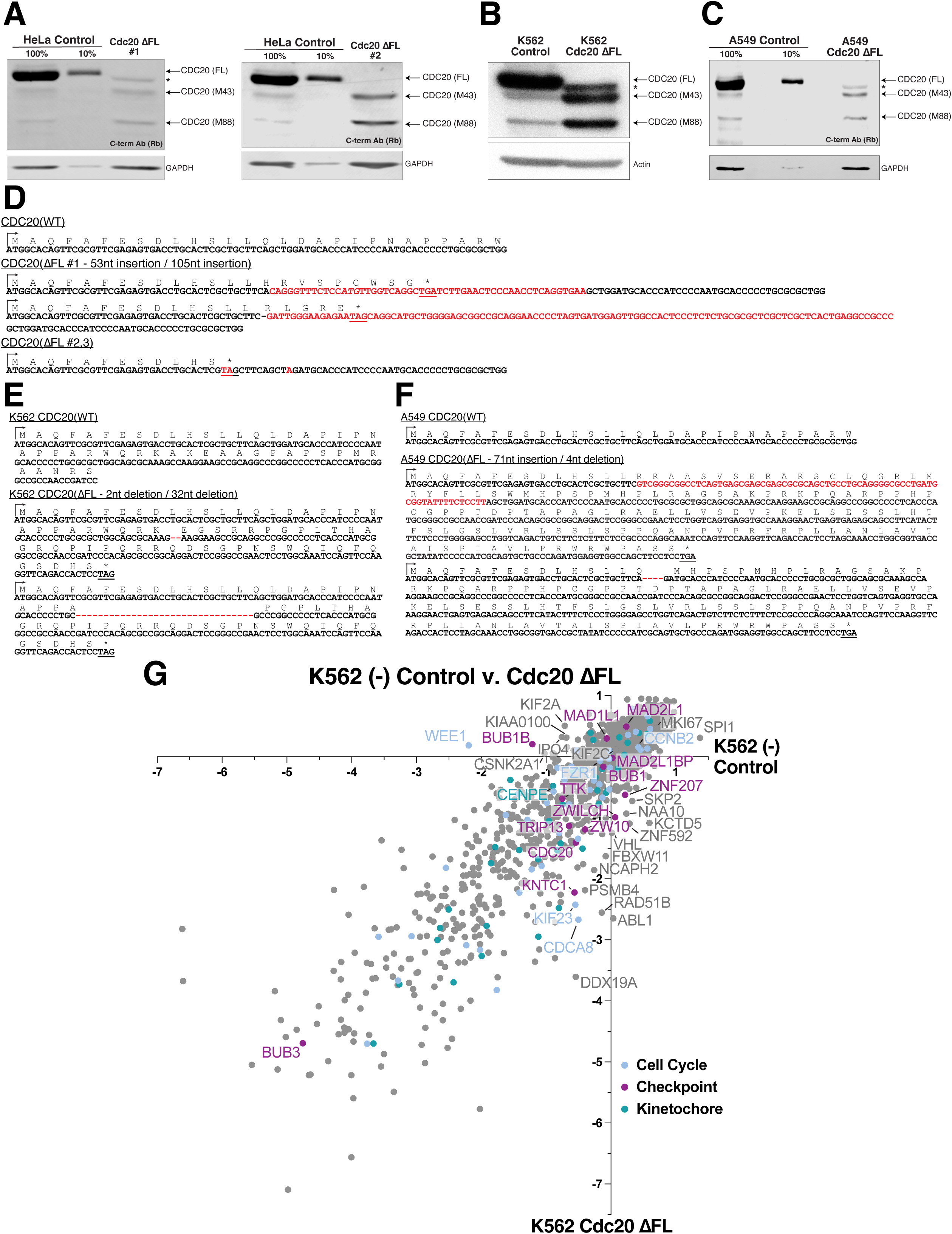
Functional genetics screen reveals differential modulators of mitotic fidelity and progression in Cdc20 ΔFL cells compared to control cells. Related to Fig. 1. **a,** Western blot analysis of endogenous Cdc20 in HeLa and Cdc20 ΔFL (Left: Clone #1, from previous study ^7^; Right: Clone #2, this study) using antibodies recognizing the C-terminus of human Cdc20 (aa 450-499). GAPDH was used as loading control. Starred is a cryptic Cdc20 protein product that likely arose due to the introduction of a predicted alternative ATG start site that connects with downstream regions of the CDC20 protein during the Cdc20 ΔFL cell line production. See **d** for more details and previous work^7^ for a detailed explanation. **b,** Western blot analysis of endogenous Cdc20 in K562 and K562 Cdc20 ΔFL cells using antibodies recognizing the C-terminus of human Cdc20 (aa 450-499). Actin was used as loading control. Starred is a cryptic Cdc20 protein product that arose due to the frameshift mutation introduced in the K562 Cdc20 ΔFL cell line connecting the predicted altORF sequence with downstream regions of the CDC20 protein. See **e** for more details and previous work^7^ for a detailed explanation. **c,** Western blot analysis of endogenous Cdc20 in A549 and A549 Cdc20 ΔFL cells using antibodies recognizing the C-terminus of human Cdc20 (aa 450-499). GAPDH was used as loading control. Starred is a cryptic Cdc20 protein product that likely arose due to the introduction of an alternative start site that connects with downstream regions of the CDC20 protein during the Cdc20 ΔFL cell line production. See **f** for more details and previous work^7^ for a detailed explanation. **d,** Sequence information for HeLa Cdc20 ΔFL (Clones #1-3) cell lines. Cdc20 ΔFL (Clone #1) contains insertions of 53 nt and 105 nt (in red) respectively after the L14 residue. The alternative ATG start site (in allele with 53 nt insertion) that connects with downstream regions of the CDC20 protein is italicized. Cdc20 ΔFL (Clones #2,3) has a 2 nt nonsense mutation at the L13 residue (CT to TA, in red) that introduces a premature stop codon, and a 1 nt missense mutation (G to A, in red) at the L16 residue that mutates the PAM site for a guide RNA sequence targeting *CDC20* Exon 1. Underlined are premature stop codons. The DNA sequence was determined by next-generation sequencing (Clone #1) and Sanger sequencing (Clones #2,3). **e,** Sequence information for K562 Cdc20 ΔFL cell line. K562 Cdc20 ΔFL contains deletion (in red) of 2 nt and 32 nt at the E32 and A26 residues, respectively. The alternative ATG start site that connects with downstream regions of the CDC20 protein is italicized. Underlined are premature stop codons. The DNA sequence was determined by Sanger sequencing. **f,** Sequence information for A549 Cdc20 ΔFL cell line containing a 71 nt insertion after L14 residue and 4 nt deletion at the Q15 residue (GCTG) in red, respectively. Underlined are premature stop codons. The DNA sequence was determined by Sanger sequencing. **g,** Scatter plot showing the CRISPR scores in K562 (-) control cells vs. K562 Cdc20 ΔFL cells.

**Fig. S2:**
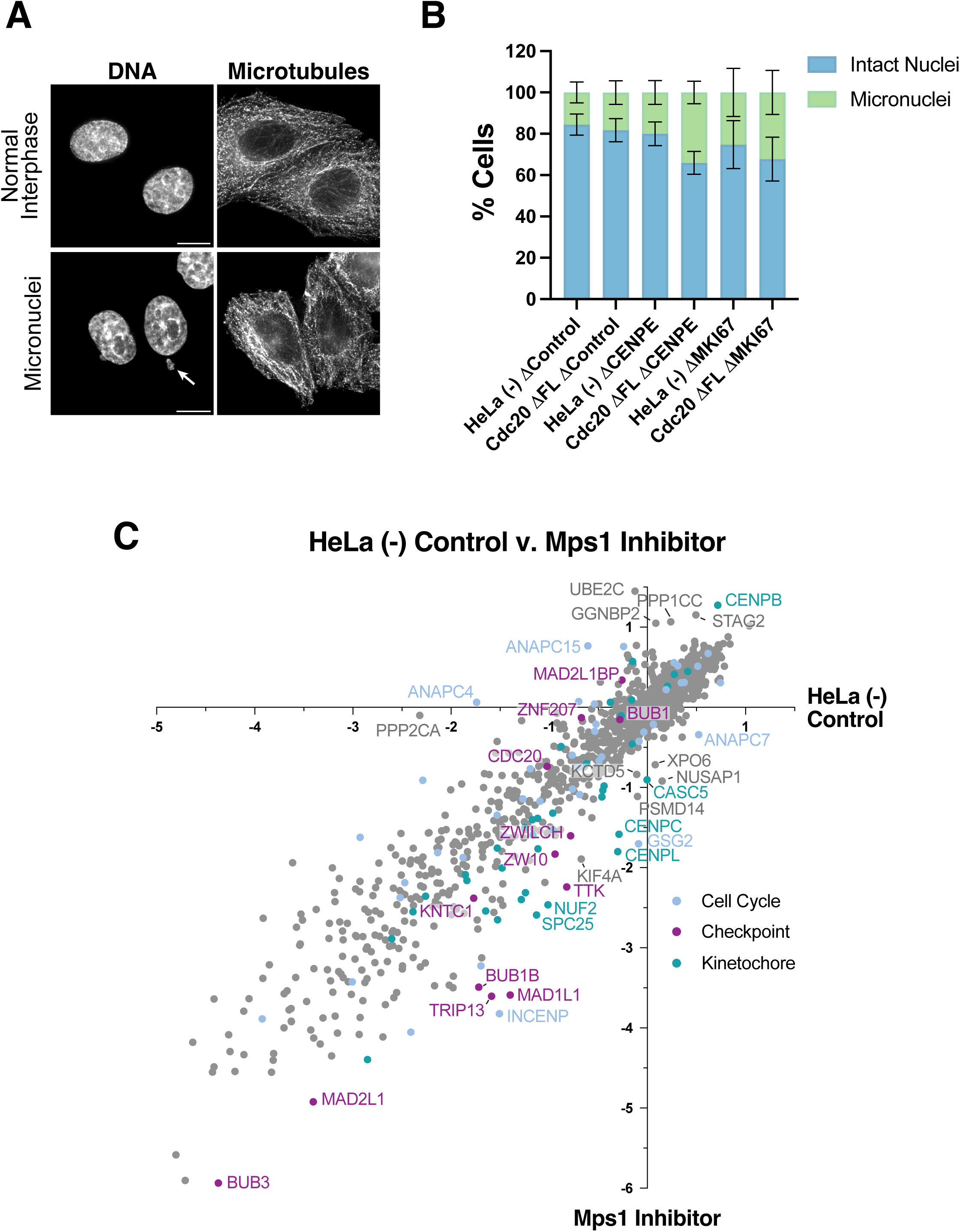
Analysis of micronuclei formation in Cdc20 ΔFL cells treated with an additional CENPE or MKI67 knockout. Related to Fig. 2. **a**, Representative deconvolved Z-projected immunofluorescence images of interphase cells from HeLa and Cdc20 ΔFL (Clone #1) cells in which CENPE or MKI67 is inducibly eliminated. Images show microtubules (DM1α) and DNA (Hoeschst). Arrowheads indicate micronuclei. Scale bars, 10 µM. **b,** Percentage of cells with micronuclei in HeLa and Cdc20 ΔFL (Clone #1) cells with a single control locus (ΔHS1), CENPE, or MKI67 knockout. *n* > 400 cells per cell line, across four experimental replicates. Mean percentage is shown, error bars indicate SD. **c,** Scatter plot showing the CRISPR scores in HeLa (-) control cells vs. Mps1 inhibitor treated HeLa cells.

**Fig. S3:**
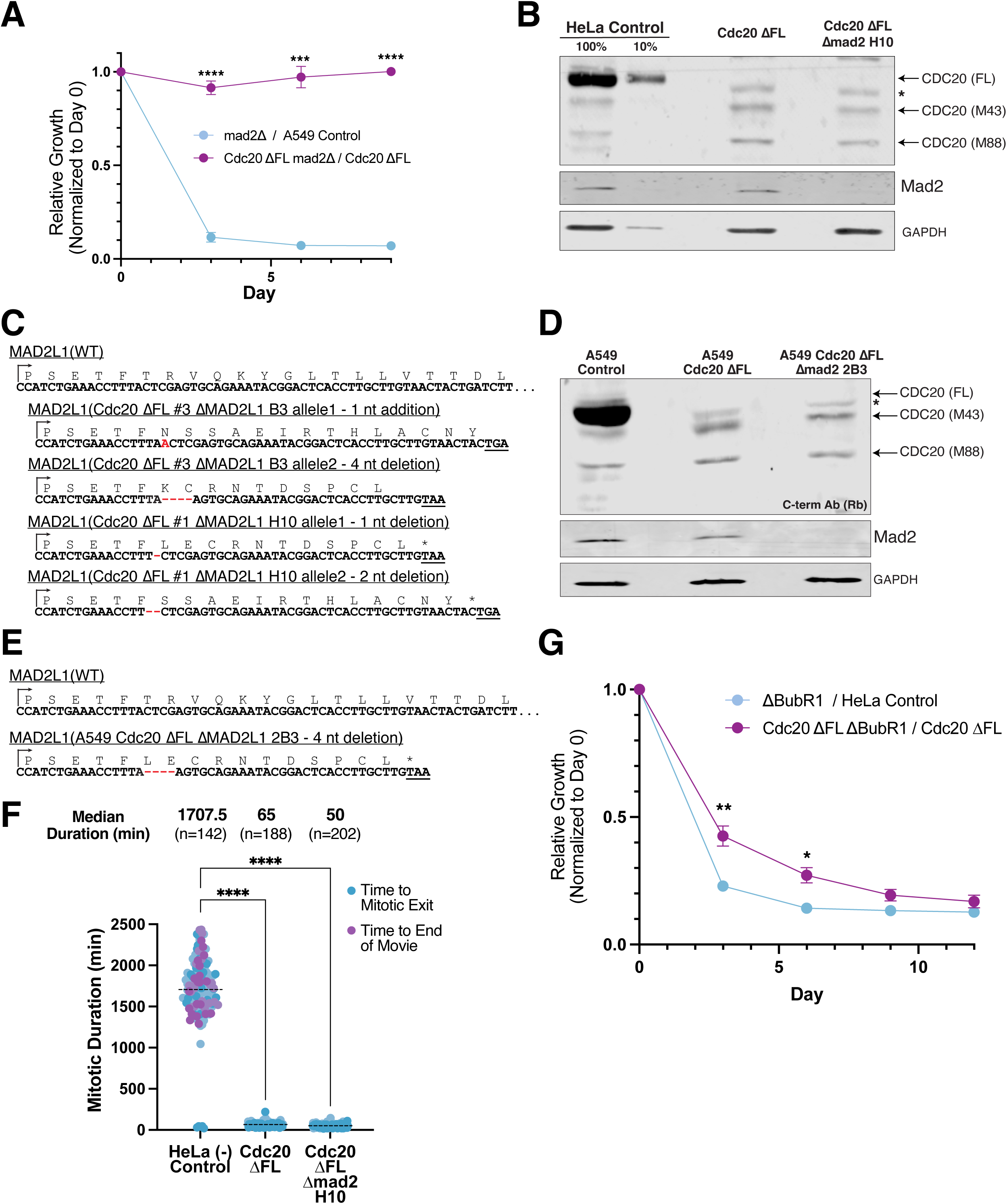
Mad2 and other Spindle Assembly Checkpoint proteins are dispensable in Cdc20 ΔFL cells. Related to Fig. 3. **a,** Competitive growth assay to test the relative fitness of inducibly eliminating Mad2 in A549 control or A549 Cdc20 ΔFL cells. Data points reflect mean ± s.e.m; *n =* 3 experimental replicates. **b,** Western blot analysis of endogenous Cdc20 and Mad2 in HeLa (-) control, Cdc20 ΔFL (Clone #1), and Cdc20 ΔFL Δmad2 H10 cells using antibodies recognizing the C-terminus of human Cdc20 (aa 450-499). Cdc20 ΔFL Δmad2 H10 cells were derived from the Cdc20 ΔFL (Clone #1) cell line (see Methods). GAPDH was used as loading control. **c,** Sequence information for the HeLa Cdc20 ΔFL Δmad2 B3 (derived from Cdc20 ΔFL Clone #3; A insertion and 4 nt deletion in red) and the HeLa Cdc20 ΔFL Δmad2 H10 (derived from Cdc20 ΔFL Clone #1; deletions of 1 and 2 nt in red) cell lines. Underlined are premature stop codons. The DNA sequence was determined by Sanger sequencing. **d,** Western blot analysis of endogenous Cdc20 and Mad2 in A549, A549 Cdc20 ΔFL, and A549 Cdc20 ΔFL Δmad2 2B3 cells using antibodies recognizing the C-terminus of human Cdc20 (aa 450-499). GAPDH was used as loading control. **e,** Sequence information for the A549 Cdc20 ΔFL Δmad2 2B3 cell line showing 4 nt deletion in both alleles in red. Underlined are premature stop codons. The DNA sequence was determined by Sanger sequencing. **f,** Mitotic arrest duration of HeLa, Cdc20 ΔFL (Clone #1), Cdc20 ΔFL Δmad2 H10 cells treated with STLC. Shown are cells entering in the first 24 hours that either exit mitosis (blue) or remain arrested (purple). Data are median across two experimental replicates; replicates are color coded. The total number of cells analyzed is indicated. **g,** Competitive growth assay to test the relative fitness of inducibly eliminating BubR1 in HeLa or Cdc20 ΔFL (Clone #1) cells. Data points reflect mean ± s.e.m.; *n* = 3 experimental replicates. For **a,g**, Student’s two-sample-*t-*tests with two-tailed distribution comparing relative ratios shown for Days 3-12. \**P*<0.05, \*\**P*<0.01, \*\*\**P*<0.001, \*\*\*\**P*<0.0001. For **f**, the Kruskal-Wallis and Dunn’s post-hoc test was performed comparing mitotic timings. \*\*\*\**P*<0.0001.

**Fig. S4:**
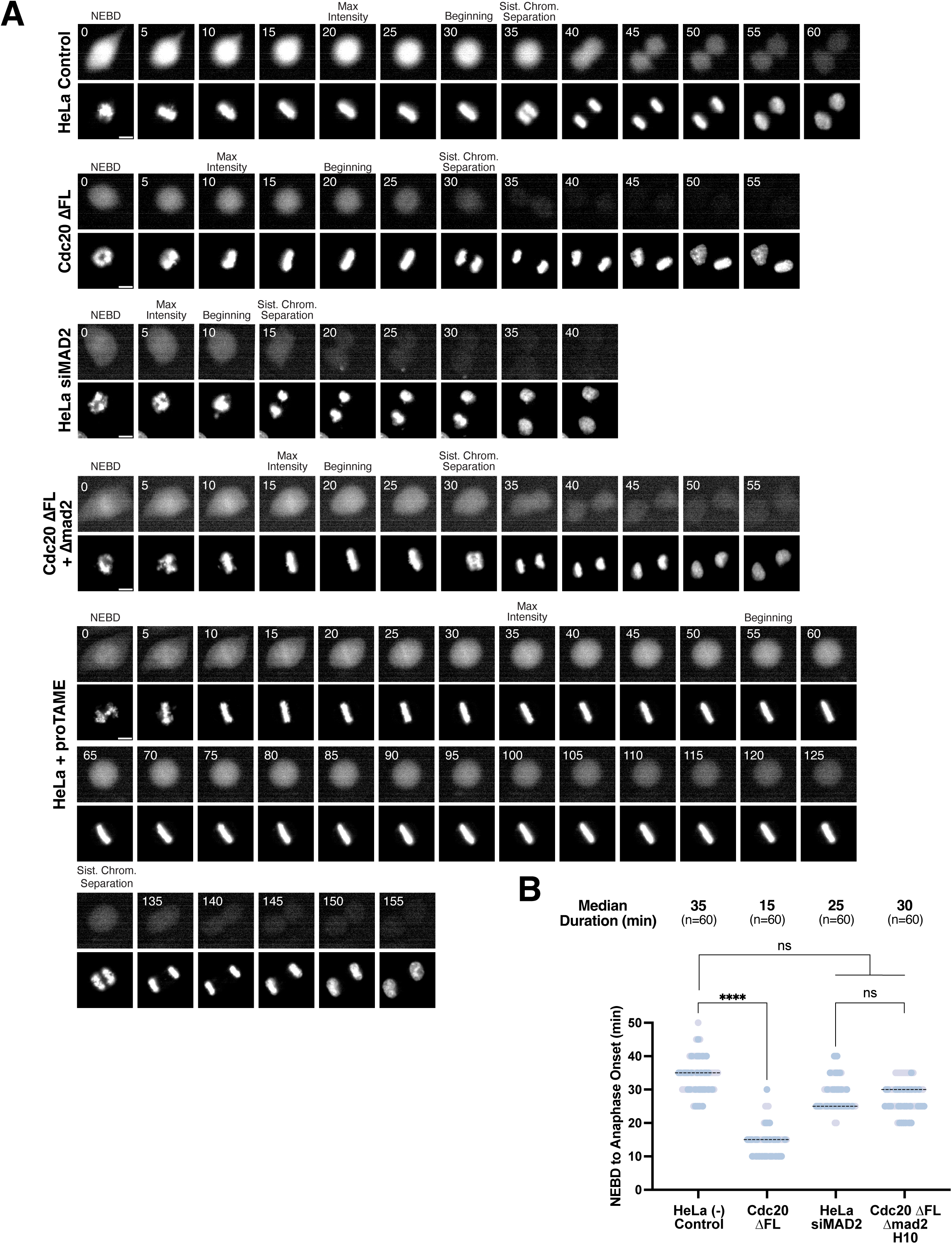
Cyclin B1 degradation kinetics and activation timing in Cdc20 ΔFL and Cdc20 ΔFL Δmad2 cells. Related to Fig. 4. **a,** Representative stills of HeLa, Cdc20 ΔFL (Clone #1), HeLa siMAD2, Cdc20 ΔFL Δmad2, and HeLa proTAME cells from NEBD (t = 0 min) to 25 minutes after sister chromatid separation taken every 5 minutes. Images show Cyclin B1 endogenously tagged with Venus fluorescence and DNA (SIR-DNA). See **4a** for definitions and further explanation. **b,** Quantification of timing from NEBD to sister chromatid separation for HeLa, Cdc20 ΔFL (Clone #1), HeLa siMAD2, Cdc20 ΔFL Δmad2 cells. Data are median across two experimental replicates; replicates are color coded. The total number of cells analyzed is indicated. The Kruskal-Wallis and Dunn’s post-hoc test was performed on the NEBD to sister chromatid separation timings. \*\*\**P*<0.001, \*\*\*\**P*<0.0001, ns>0.9999.

**Fig. S5:**
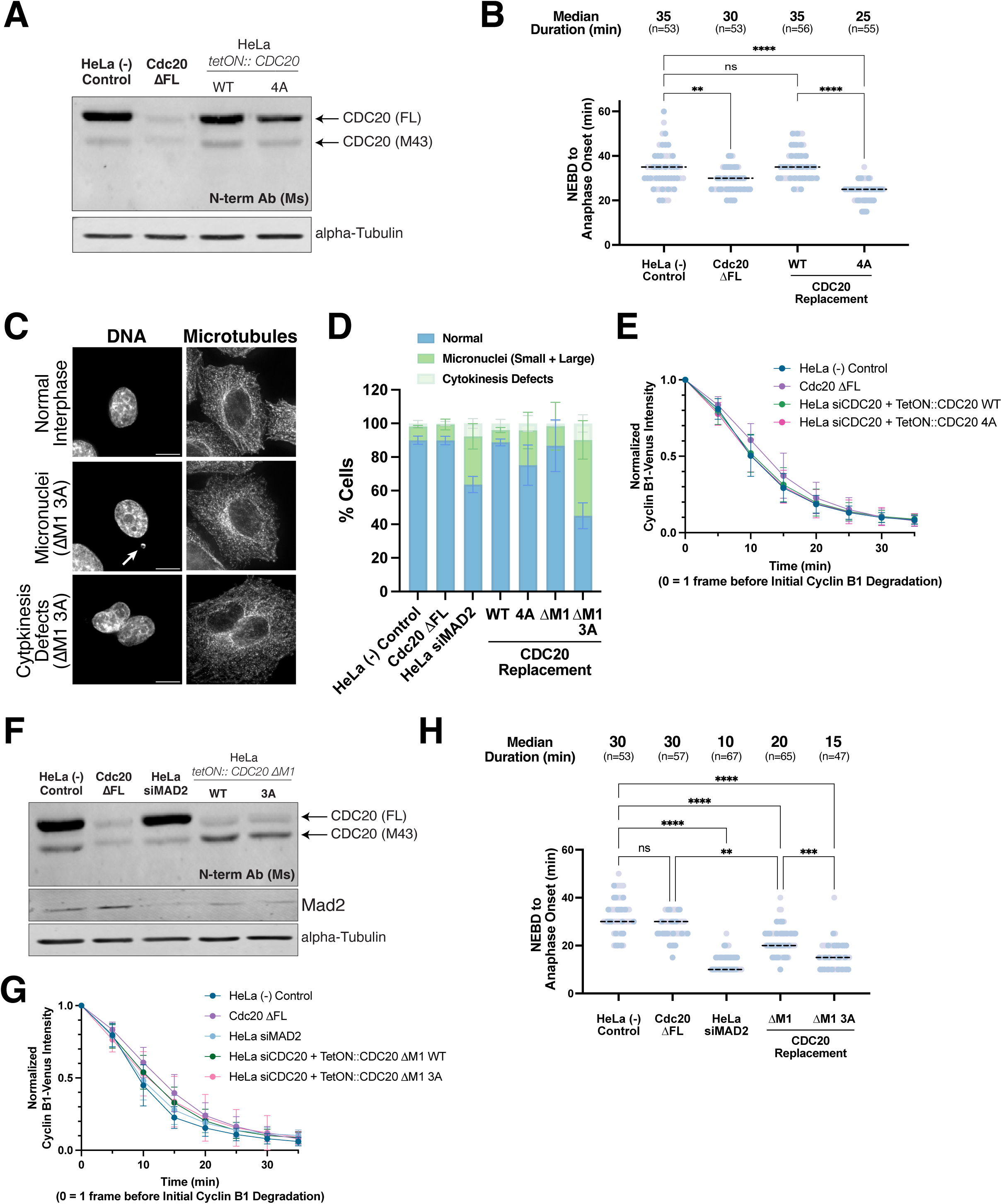
Cdk1-mediated phosphorylation of Cdc20 regulates the timing of APC/C activation. Related to Fig. 5. **a,** Western blot analysis of Cdc20 in HeLa cells, Cdc20 ΔFL (Clone #1) cells, and cells in which endogenous CDC20 protein is replaced with either WT siRNA-resistant *CDC20* cDNA (WT) or a mutant with four Cdk1-mediated phosphorylation sites (S41, T55, T59, T70) mutated to alanine (4A) using antibodies recognizing the N-terminus of human Cdc20 (aa 1-175). α-Tubulin was used as loading control. **b,** Quantification of timing from NEBD to sister chromatid separation for cell lines described in **a**. Data are median across two experimental replicates; replicates are color coded. The total number of cells analyzed is indicated. **c,** Representative deconvolved Z-projected immunofluorescence images of interphase cells from cell lines indicated in **d**. Images show microtubules (DM1α) and DNA (Hoeschst). Arrowheads indicate micronuclei. Scale bars, 10 µM. **d,** Percentage of cells with micronuclei in cell lines indicated. *n* > 150 cells per cell line, across three experimental replicates. Percent mean is shown, error bars indicate SD. **e,** Cyclin B1 degradation curves for cell lines indicated in **a** during anaphase. Time t = 0 min is defined as 5 minutes before onset of cyclin B1 degradation (onset of cyclin B1 degradation is defined as a decrease in CCNB1-Venus intensity >10%), and all time points (every 5 minutes for 35 minutes) are normalized to the CCNB1-Venus fluorescence intensity at t = 0. Data points reflect mean ± s.e.m.; *n* = ∼60 cells across two experimental replicates (exact number of cells seen in **b**). **f,** Western blot analysis of Cdc20 in HeLa cells, Cdc20 ΔFL (Clone #1) cells, HeLa siMAD2 cells, and cells in which endogenous CDC20 protein is replaced with a mutant disrupting the Met1 start site (ΔM1) or a mutant disrupting the Met1 start site with also three Cdk1-mediated phosphorylation sites mutated (T55, T59, T70) to alanine (ΔM1 3A) using antibodies recognizing the N-terminus of human Cdc20 (aa 1-175). α-Tubulin was used as loading control. **g,** Cyclin B1 degradation curves for cell lines indicated in **f** during anaphase. See **e** for more details (exact number of cells seen in **h**). **h,** Quantification of timing from NEBD to sister chromatid separation for cell lines described in **f**. Data are median across two experimental replicates; replicates are color coded. The total number of cells analyzed is indicated. For **b,h**, the Kruskal-Wallis and Dunn’s post-hoc test was performed on the NEBD to sister chromatid separation timings. \*\**P*<0.01,\*\*\**P*<0.001, \*\*\*\**P*<0.0001, ns>0.9999.

**Fig. S6:**
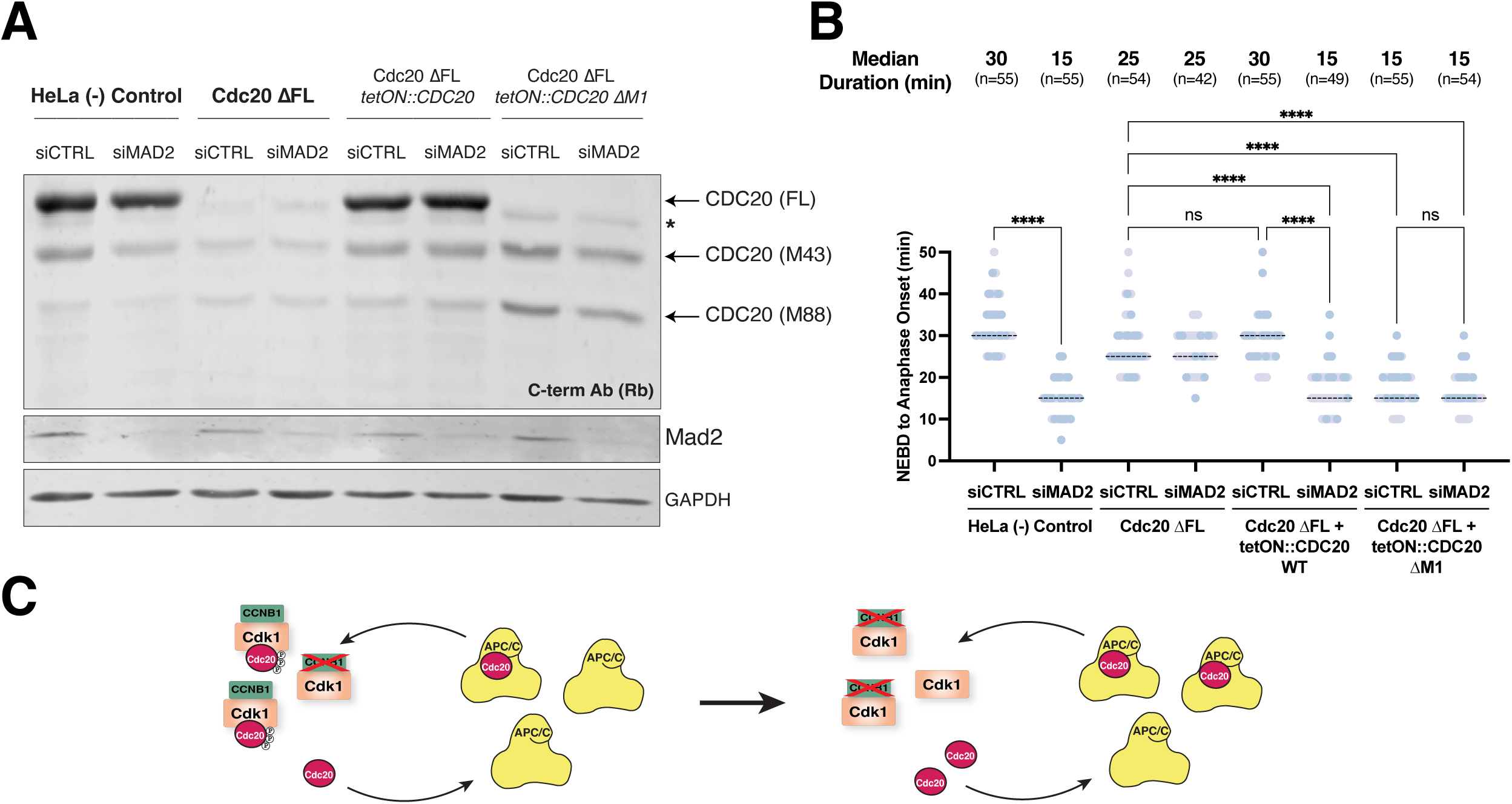
Modulating Cdc20 protein levels in Cdc20 ΔFL cells affects the timing of sister chromatid separation. Related to Fig. 6. **a,** Western blot analysis of Cdc20 in HeLa cells, Cdc20 ΔFL (Clone #1) cells, HeLa siMAD2 cells, and cells treated with 50 ng/mL doxycycline to induce expression of indicated *CDC20* constructs and RNAi against Mad2 or a non-targeting control pool using antibodies recognizing the C-terminus of human Cdc20 (aa 450-499). GAPDH was used as loading control. **b,** Quantification of timing from NEBD to sister chromatid separation for cell lines described in **a**. Data are median across two experimental replicates; replicates are color coded. The total number of cells analyzed is indicated. The Kruskal-Wallis and Dunn’s post-hoc test was performed on the NEBD to sister chromatid separation timings. \*\*\*\**P*<0.0001, ns > 0.9999. **c,** Model of feedback loop involving Cdc20 and CDK activity that regulates the timing of APC/C activation.

**Supplementary Table 1.** CRISPR/Cas9 screen sgRNA sequences, counts, and gene scores (see attached Excel file)

**Supplementary Table 2.** Antibodies used in study.

| **Antibody** | **Source** | **Raised in** | **Dilution/Use** |
| --- | --- | --- | --- |
| Cdc20 C-terminus (aa 450-499) | Abcam ab26483, lot 1052555-13 | Rabbit polyclonal | 1:500 (WB) |
| Cdc20 N-terminus (1-175) | Santa Cruz Biotech sc13162, lot I1820/I0321/G1923 | Mouse clone E-7 | 1:200 (WB) |
| Mad2 | Abcam ab9777, lot 1107907-2; GR2167-51 | Rabbit polyclonal | 1:5000 (WB) |
| GAPDH | Cell Signaling Tech #2118, lot H1224 | Rabbit clone 14C10 | 1:1000 (WB) |
| alpha tubulin (DM1α) | Cell Signaling 3873 | Mouse monoclonal | 1:1000 (IF) |
| alpha tubulin | Abcam ab52866, lot 1011212-35 | Rabbit monoclonal | 1:5000 (WB) |
| β-Actin (HRP-conjugated) | Santa Cruz Biotech sc-47778, lot G0319 | Mouse clone C4 | 1:20,000 (WB) |

WB = Western blot, IF = immunofluorescence

**Supplementary Table 3.**
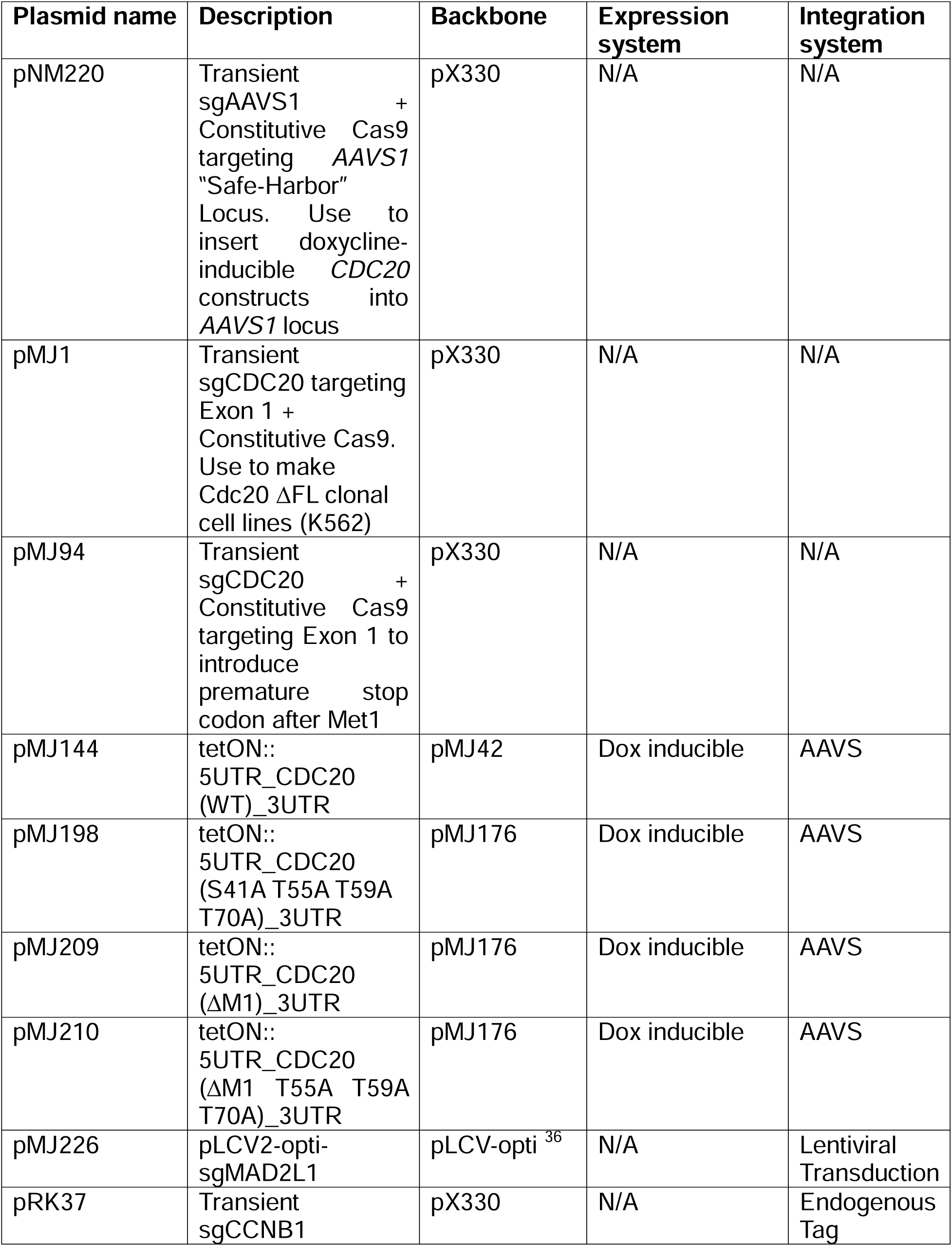
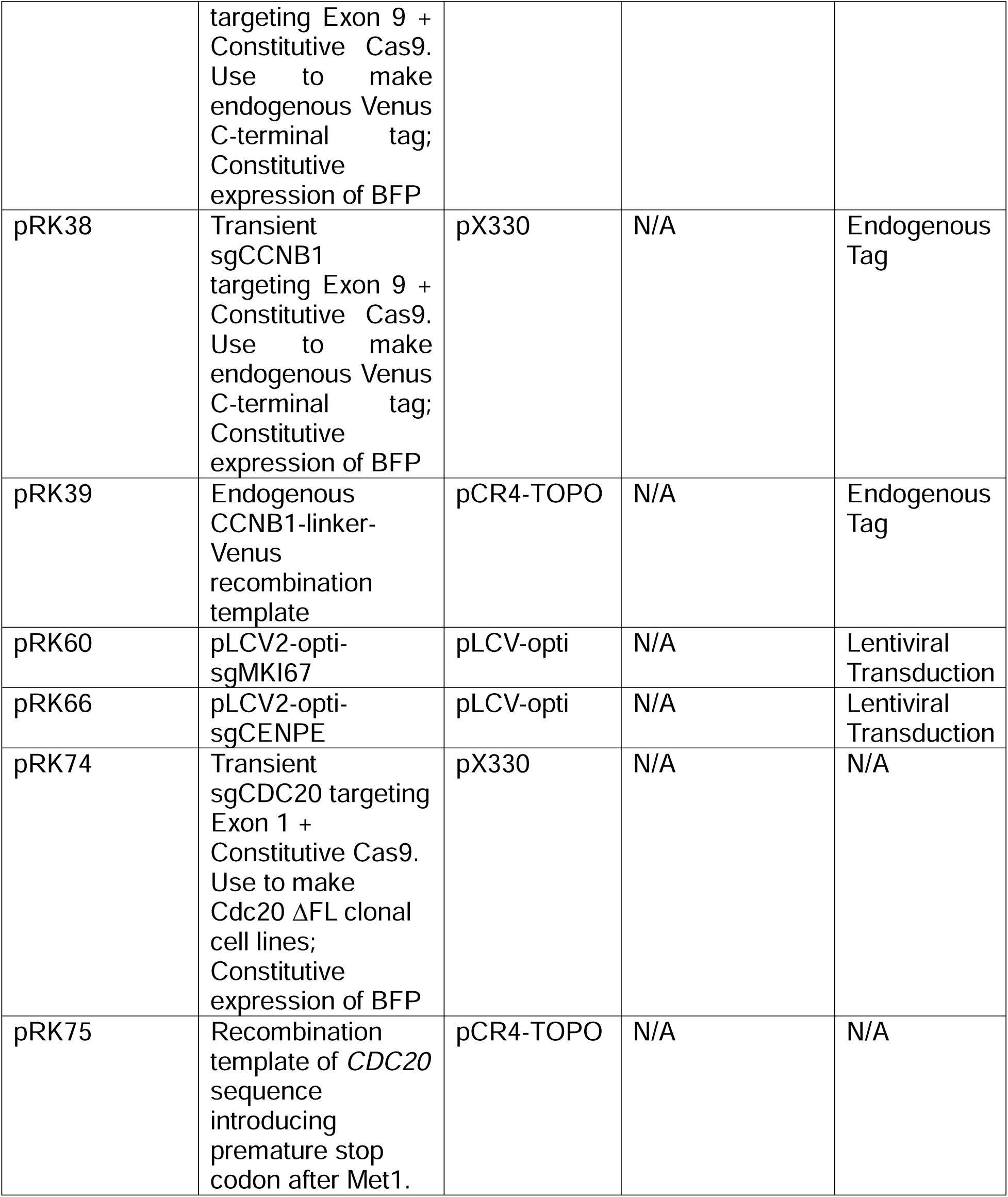
Plasmids used in this study.

**Supplementary Table 4.**
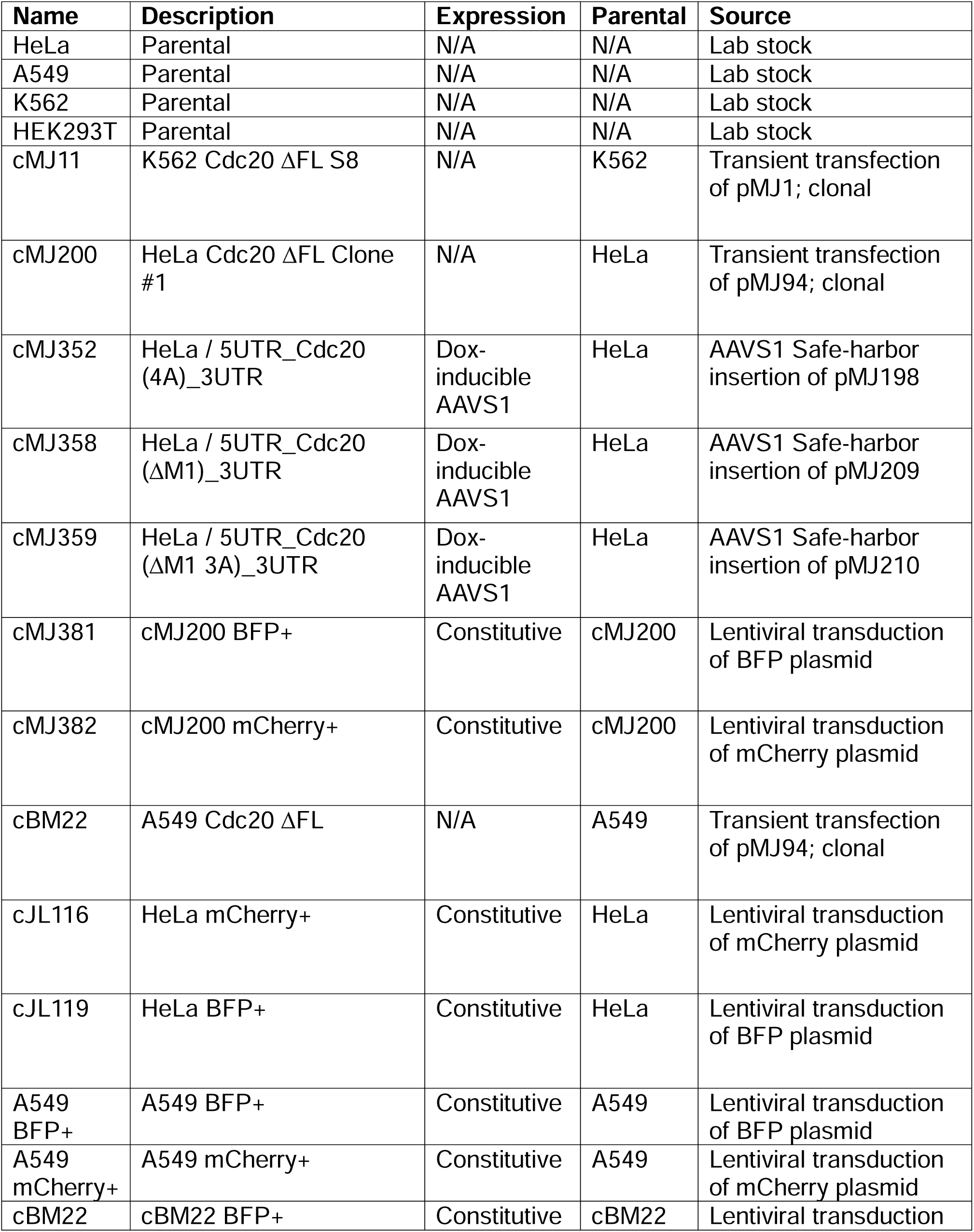
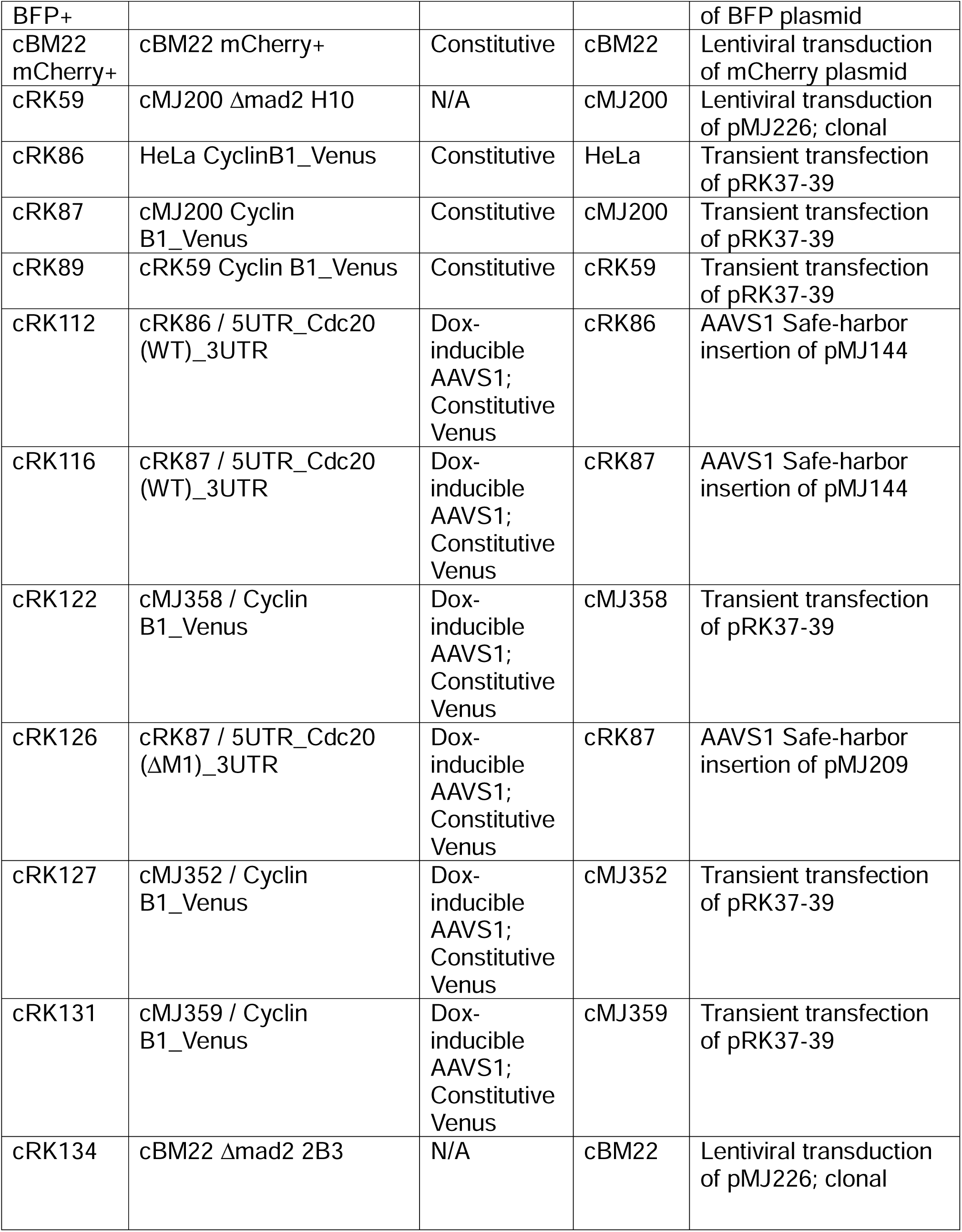
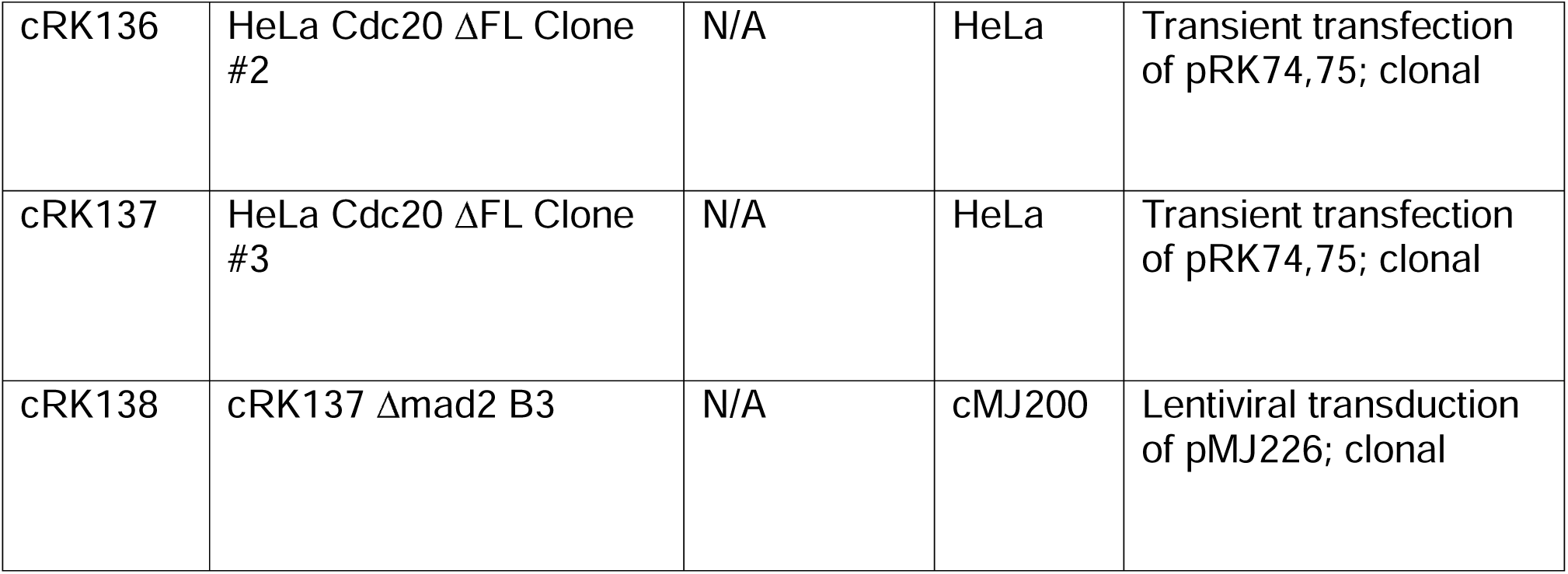
Cell lines used in this study.

